# TNF-TNFR Signaling Modality Conserved Across Invertebrate

**DOI:** 10.1101/2024.07.27.603686

**Authors:** Manoj K. Govindan, Kaushiki K, Moushmi Goswami, Namita Menon, Anmol Singh, Subhashini Srinivasan

## Abstract

In humans, the signaling mechanisms of the 19 paralogs of the tumor necrosis factor superfamily (TNFSF) and the 29 receptor paralogs of the tumor necrosis factor receptor superfamily (TNFRSF) are extensively characterized because of their therapeutic relevance. The functional expansion of TNFSF in vertebrates from a single ancestral gene through successive duplication events is also well established. However, apart from the first identification of a TNFSF homolog, Eiger (dmEiger), in *Drosophila melanogaster* in 2002, together with its receptor homologs Wengen (dmWgn) and Grindelwald (dmGrnd), this signaling system has remained largely unexplored in invertebrates. More recently, the implication of an Eiger homolog in *Plasmodium* resistance in malaria vectors has further highlighted the need for a systematic investigation of this pathway in lower invertebrates.

Structural comparison of the dmEiger-dmGrnd complex with the canonical 3:3 ligand-receptor configuration observed in human TNFSF-TNFRSF signaling suggests either conservation of this signaling modality since before the bilaterian split or convergent evolution of a similar architecture in both branches. The recent explosion in high-quality proteomes spanning diverse phyla, together with advances in protein-complex prediction using AlphaFold-multimer, now enables large-scale exploration of ligand-receptor evolution across invertebrates.

Here, we analyzed 148 near-complete proteomes spanning major invertebrate phyla and identified 290 TNFSF, 336 wengen (wgn), and 115 grindelwald (grnd) homologs, including homologs from lower invertebrates. Structural characterization of 140 selected complexes using AlphaFold and AlphaFold-multimer revealed several key findings: (i) TNFSF and TNFRSF homologs are present in majority of the phyla under invertebrates (ii) the canonical 3:3 ligand-receptor signaling configuration is conserved across invertebrates; (iii) orthologs of 25 out of the 26 genes implicated in TNF signaling pathways are present in lower invertebrates; and (iv) signaling through grnd-like receptors containing a single cysteine-rich domain with CXXCXXXC signature is the predominant signaling mode in invertebrates and becomes highly prevalent in Arthropoda.

We also elaborate a hypothesize on the evolutionary trajectories toward a genetically parsimonious signaling by this complex system before functional expansion in vertebrates and species diversification in Arthropoda.

## INTRODUCTION

Tumor necrosis factor superfamily (TNFSF)-tumor necrosis factor receptor superfamily (TNFRSF) signaling plays a central role in vertebrate immunity and inflammation, and inhibitors targeting these pathways are widely used in the treatment of autoimmune disorders (Sonar and Lal 2015). Viruses have evolved multiple strategies to interfere with this pathway. For example, vaccinia virus encodes a TNFRSF homolog capable of blocking TNF-α signaling in humans (Tian et al. 2012). In another example, viral capsid proteins may have evolved from TNFSF-like proteins, or vice versa, through ancient evolutionary exchange between viruses and cellular organisms (Krupovic and Koonin 2017). The large variation in the number of TNFSF ligands and receptors across vertebrates raises important questions regarding the evolutionary mechanisms that shaped the expansion of this signaling system.

The first evolutionary model describing co-evolution of the TNFSF and TNFRSF families in vertebrates was proposed in 2003 (Collette et al. 2003). Using eight osteichthyan species, the authors suggested that all vertebrate TNFSF genes originated from a single ancestral gene that underwent tandem duplication, generating four genes distributed across two chromosomes. Subsequent whole-genome duplication events (WGD1 and WGD2) expanded this repertoire to fifteen genes. This framework was later refined using homologs from additional clades (Marín 2020). More recently, a comprehensive evolutionary analysis incorporating cyclostomes proposed a simplified model of TNFSF evolution in early vertebrates (Marín 2025). Figures 14 and 15 from this report provide a detailed account of duplication and gene-loss events that ultimately produced the 19 TNFSF paralogs found in humans. Together, these studies strongly support the view that vertebrate TNFSF members emerged through sequential duplication events followed by selective gene loss.

Experimentally determined TNFSF-TNFRSF complexes reveal a conserved 3:3 homotrimeric signaling architecture in both arthropods and vertebrates. This arrangement is observed in the dmEiger-dmGrnd complex from *Drosophila melanogaster* (PDB: 6zt0, 6i50) and in 15 of the 19 human TNFSF paralogs complexed with their respective receptors (PDB: 3alq, 4v46, 1xu1, 1d0g, 3qd6, 6mkz/6bwv, 3qbq, 4msv, 7kx0, 1oqd, 2hey, 1rj7, 4en0, 2rjk, 7khd, 4ht1). The conservation of this architecture suggests that homotrimeric signaling predates the bilaterian split and may represent an evolutionarily parsimonious solution for ligand-receptor signaling.

The functional importance of homotrimerization is well established in vertebrates. For example, membrane anchoring is essential for the activity of CD40L; unanchored forms lose signaling activity because of impaired trimerization. Restoration of trimerization through engineered coiled-coil domains rescues activity (Fanslow et al. 1994; Gurunathan et al. 1998). Likewise, disruption of TNF-α trimerization has been explored as a therapeutic strategy to inhibit signaling (PDB: 7ta6).

Despite extensive characterization in vertebrates, the TNFSF-TNFRSF system remains poorly explored in invertebrates. Among invertebrate homologs, Eiger was first identified in *Drosophila melanogaster* (Igaki et al. 2002), together with two receptor homologs, Wengen (dmWgn) and Grindelwald (dmGrnd) (Kanda et al. 2002). Subsequent studies demonstrated that N-terminal glycosylation of dmGrnd modulates JNK signaling by reducing dmGrnd-dmEiger binding affinity (Palmerini et al. 2021). More recently, homologs of Eiger, wengen, and grnd were identified in the malaria vector *Anopheles stephensi*, where Eiger localizes within an inversion region unique to strains with low vectorial capacity, suggesting a potential role in parasite resistance (Srinivasan et al. 2022).

Evidence for TNFSF-TNFRSF homologs in invertebrate phyla has gradually emerged across multiple phyla. In Mollusca, the first TNFSF homolog was characterized in *Haliotis discus*, where expression was significantly upregulated in gill tissues exposed to pathogens (De Zoysa et al. 2009). The first TNFRSF homolog in scallop (*Chlamys farreri*) was similarly found to be induced following exposure to gram-negative bacteria (Li et al. 2009). Additional studies in *Anodonta woodiana* and *Pinctada fucata martensii* implicated TNFRSF homologs in immune defense pathways (Li et al. 2009; Wu et al. 2020). A large-scale expansion of TNFSF-TNFRSF homologs, reminiscent of vertebrate diversification, has also been reported in Pacific oyster (*Crassostrea gigas*) (Zhang et al. 2015). In Platyhelminthes, five TNFSF homologs from non-parasitic species and thirty-one TNFRSF homologs have been reported (Bertevello et al. 2020).

In Cnidaria, TNFSF-TNFRSF homologs appear to predate the vertebrate-invertebrate split and have been implicated in heat-stress responses (Quistad and Traylor-Knowles 2016). Multiple TNFRSF homologs identified in *Acropora digitifera* can cross-react with human TNF-α, suggesting deep functional conservation (Quistad et al. 2014). Homologs of TNFSF and TNFRSF have also been identified in *Hydra* (Steichele et al. 2021). Furthermore, heat-resistant coral reefs display upregulation of TNFRSF homologs and associated pathway genes (Barshis et al. 2013). In Porifera, the first TNFSF-TNFRSF homologs were identified in the marine sponge *Chondrosia reniformis* (Pozzolini et al. 2016), while immune cross-reactivity against antibodies targeting human TNF-α suggests the existence of TNF-like molecules in additional sponge species.

Here, leveraging the rapid expansion of high-quality proteomes across diverse phyla and recent advances in protein-complex prediction using AlphaFold-multimer, we investigate the evolution of TNFSF-TNFRSF signaling across Metazoa, with a particular focus on lower invertebrates.

## RESULTS

We scanned 148 completed genomes/proteomes of species representing all major invertebrate phyla to investigate the presence, absence, and paralog expansion of members of the TNFSF (Eiger) and TNFRSF (grnd/wgn) families across all phyla under invertebrates. As described under the Materials section these species were strategically chosen to minimize false negatives during mining.

### TNFSF/TNFRSF homologs/paralogs mined across invertebrates

Figure 1 summarizes the results from mining the complete proteomes of the 148 species. In total, we identified 290 Eiger homologs, 336 wgn homologs, and 115 grnd homologs. The link for the detailed annotations, including sequences and UniProt accessions for homologs identified in each species and phylum are provided in the Data Availability section. The cladogram in Figure 1a illustrates the number of proteomes analyzed per phylum, while the histogram in Figure 1b shows the distribution of paralog counts per species across phyla, with Eiger shown in red, wgn in blue, and grnd in green. Supplementary Table S4 and Supplementary Figure S6 provide detailed presence/absence patterns across all species examined. We observed that TNSFSF and TNFRSF homologs are missing under Nematode, Tardigrada, and Chelicerata branch under Arthropoda.

**Figure 1:**
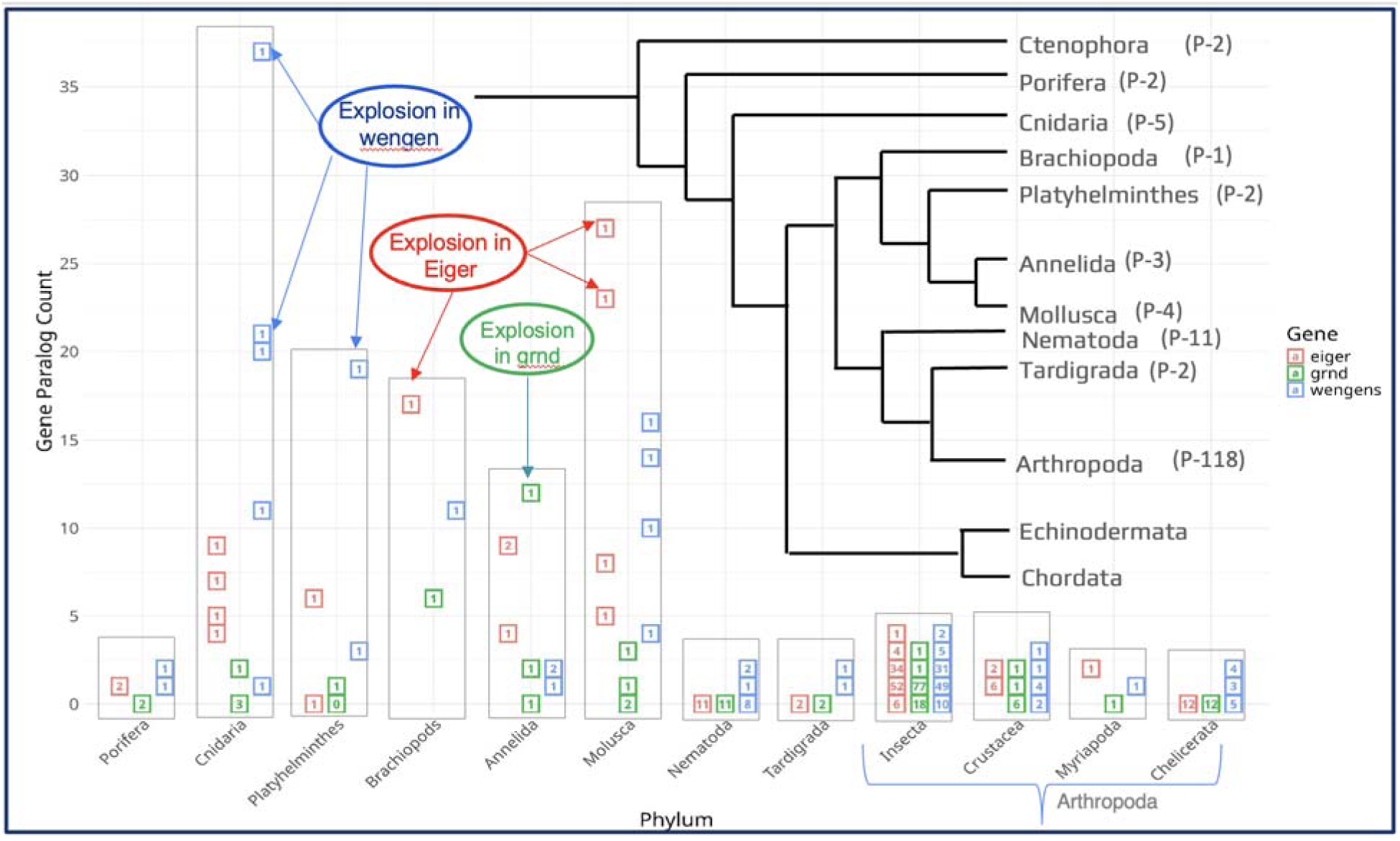
The cladogram shows the number of proteomes studies (P), while the histogram shows the number of paralogs for each species with TNFSF (red box), wgn (blue box) and grnd (green box) indicated with inset numbers representing number of species against number of paralogs on y-axis.

No clear correlation was observed between the number of Eiger homologs and the number of receptor homologs (see arrows in Figure 1b) within species. The largest expansion of Eiger paralogs was observed in Mollusca, where one species contained 27 paralogs, followed by Cnidaria with up to 9 paralogs (red arrows in Figure 1b). The highest expansion of grnd homologs occurred in Annelida, with one species containing 12 grnd paralogs (green arrows in Figure 1b). In contrast, wgn homologs showed extensive expansion in several phyla, with maxima of 37 paralogs in Cnidaria, 19 in Platyhelminthes, and 16 in Mollusca (blue arrows in Figure 1b). Notably, wgn homologs were detected in most species examined, suggesting that wgn-like receptors may predate the emergence of canonical TNFSF-TNFRSF signaling and could have existed in ancestral lineages lacking classical ligand-mediated signaling systems, as previously suggested (Barrio-Hernandez et al., 2023). Among the 118 arthropod species analyzed, at least 52 species encoded one Eiger homolog, 77 encoded one grnd homolog, and 49 encoded one wgn homolog (Supplementary Table S4).

### Insights into Eiger and grnd paralog expansion

Figure 2a presents the phylogenetic tree of all 191 TNFSF orthologs/paralogs identified across species included in this study with the exception of a single representative from Arthropoda and including all human TNFSF paralogs, which are highlighted in red. A high-resolution version of the tree is provided in Supplementary File 2. Unlike conventional phylogenetic analyses based on a single orthologous gene, this analysis includes both orthologs and paralogs because reliable ortholog assignment across deeply divergent invertebrate lineages is not feasible for members of TNFSF and TNFRSF family. Nevertheless, in many instances the tree reveals patterns of function expansion before or after speciation. Consequently, the major clades in Figure 2a were assigned primarily based on the seven major branches emerging from the tree, without strict reliance on branch confidence values. To aid interpretation, homologs from different phyla are color coded, providing independent support for the observed clustering patterns.

**Figure 2.**
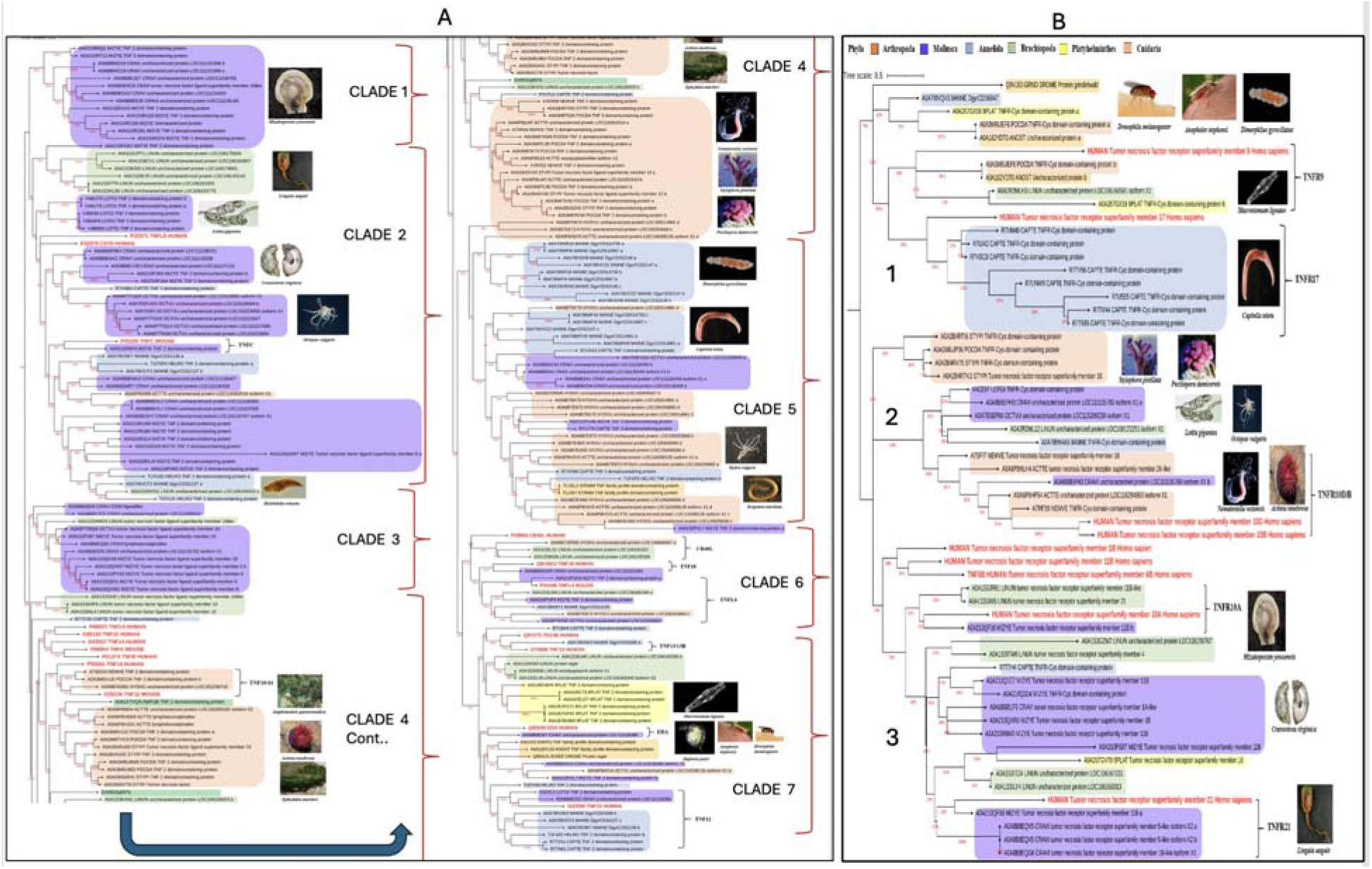
(A) Phylogenetic tree of 191 TNFSF/Eiger homologs and paralogs identified from invertebrates together with human TNFSF members, including only a few representatives from Arthropoda. (B) Phylogenetic tree of TNFRSF/grnd homologs identified across the invertebrate species analyzed in this study, together with human homologs and selected representatives from Arthropoda. Enlarged high-resolution versions of both phylogenetic trees are provided in Supplementary Files 2 and 3. Phyla are color coded as follows: Arthropoda (orange), Mollusca (purple), Brachiopoda (green), Annelida (blue), Cnidaria (pink), and Platyhelminthes (yellow).

Clades 1, 2, and 3 are dominated by homologs/paralogs from Mollusca (purple), with branch confidence values of 82%, 32%, and 46%, respectively, while Clade 4, supported by 56% confidence, is enriched for homologs from Cnidaria (pink), suggesting function expansion within each phyla. Although Clade 2 exhibits relatively low overall confidence, it predominantly contains molluscan homologs, and several internal sub-clades displaying strong support consistent with known species relationships inferred from COX-1 phylogeny (Tuliuan et al., 2025). For example, TNFSF paralogs from *Crassostrea virginia* cluster together with homologs from the closely related species *Mizuhopecten yessoensis*, suggesting that paralog expansion preceded species divergence in this lineage. In contrast, the six TNFSF paralogs identified in *Octopus vulgaris* form a highly supported monophyletic cluster within Clade 2 (100% confidence), is consistent with lineage-specific paralog expansion occurring after speciation.

Several clades contain human TNFSF representatives. However, many human paralogs form a distinct cluster under Clade 4 supporting substantial functional expansion of the TNFSF family during vertebrate evolution after the bilaterial split. The TNFSF homolog from *Drosophila melanogaster*, the sole arthropod representative included in Figure 2a, clusters with the human *EDA* gene within Clade 7 with 84% confidence.

Figure 2b shows the phylogenetic tree of all grnd homologs identified in this study including one member from dmGrnd as the sole arthropod representative. A striking expansion of grnd homologs was observed in *Capitella teleta*, where multiple paralogs cluster together within Clade 1 with 89% confidence, indicating lineage-specific duplication within this species. Similarly, grnd homologs from the molluscan species *Crassostrea virginia* and *Mizuhopecten yessoensis* cluster together within Clade 3 with 74% confidence, consistent with their close evolutionary relationship (Tuliuan et al., 2025). The single grnd homolog from *Drosophila melanogaster* clusters within Clade 1.

### Insights from unusual In-frame paralogs

Among the 290 Eiger and 336 wgn homologs identified, several species contain in-frame paralogs within the same gene, as illustrated in Figure 3 and Supplementary Figures S3 and S4. Particularly striking examples were observed in *Hydra vulgaris* (Cnidaria), where the genes A0A8B7DK70, A0A8B7E4M2, and A0A8B7E873 contain 9, 4, and 9 TNF homology domains (THDs), respectively, suggesting extreme amplification through serial domain duplication. Notably, in the AlphaFold predicted structure from UNiProt the nine THDs in A0A8B7E873 (Figure 3a) are arranged in a manner resembling a “trimer of trimers.”

**Figure 3:**
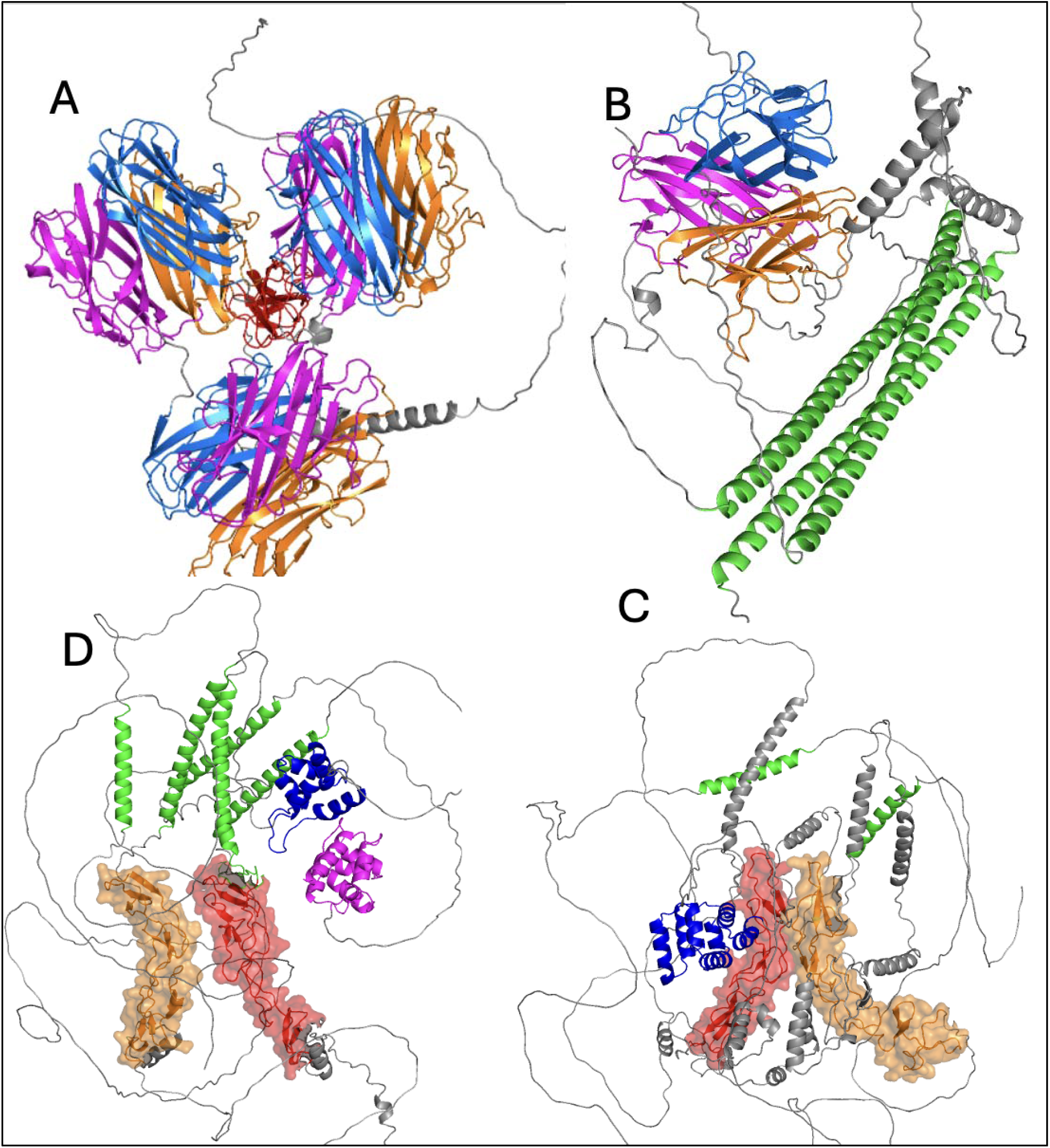
Examples of AlphaFold-predicted structures displaying complex topologies. A) An Eiger homolog from Hydra vulgaris (Cnidaria), A0A8B7E873, containing nine THDs arranged as a trimer of trimers. B) An Eiger homolog from Darwinula stevensoni, A0A7R9A6L1, containing three THD domains within a single gene. C-D) Two wengen homologs from Pocillopora damicornis (Cnidaria), A0A3M6UTV1 and A0A3M6TZR3, illustrating full-length in-frame paralogs generated through internal duplication events.

Similarly, the wgn receptor homologs A0A3M6UTV1 and A0A3M6TZR3 from *Pocillopora damicornis* (Cnidaria) each contain two distinct receptor-binding regions composed of multiple cysteine-rich domains (CRDs), separated by transmembrane (TM) regions, as shown in Figures 3c and 3d. Perhaps, these unusual ligand and receptor architectures within the TNF-TNFR system resemble evolutionary snapshots of ancient duplication events that may have been retained in extant organisms under specific selective pressures. If so, these observations underscore the extraordinary genetic burden associated with the functional expansion of TNFSF and TNFRSF paralogs through duplication.

The evolutionary and genetic burden associated with maintaining such complex ligand architectures for signaling through a single receptor system cannot be overstated, especially when contrasted with the far more parsimonious mechanism observed in humans and most arthropods, where signaling occurs through homotrimerization of three independent copies of monomers, each encoded by a separate gene containing a single THD.

### A single instance of signaling via heterotrimerization of TNFSF: A0A7R9A6L1

In *Darwinula stevensoni* (Arthropoda), a single gene, A0A7R9A6L1 (dsEiger), contains three tandem TNFSF paralogs arranged in-frame, as illustrated in Figure 3b. Each THD is associated with an obligatory N-terminal transmembrane region, a defining characteristic of the TNFSF superfamily, suggesting that this architecture likely arose through in-frame triplication events. Notably, this species encodes only a single grnd receptor homolog (dsGrnd). The three identical copies of the grnd cysteine-rich domain (CRD) engage the three ligand interfaces formed by homotrimerization of three TNFSF monomers. If dsEiger signals through heterotrimerization of its three distinct THDs using the single dsGrnd receptor, then the same dsGrnd CRD must recognize three structurally distinct ligand interfaces generated within the heterotrimeric assembly. It is also possible that the terminal THD of dsEiger is capable of independent homotrimerization and signaling.

Supporting these possibilities, the AlphaFold-predicted structure of dsEiger available in UniProt, shown in Figure 3b, indicates that the three THDs interact spontaneously as a heterotrimer even in the absence of dsGrnd. To determine whether the three resulting ligand interfaces are structurally compatible with receptor engagement, we modeled a ligand-receptor complex using AlphaFold-Multimer (AF-M). The model included the amino acid sequences corresponding to the three distinct THDs of A0A7R9A6L1 together with three identical copies of dsGrnd. The predicted complex, shown in Figure 4d, satisfied all structural and energetic criteria established from experimentally determined crystal structures (Table 1, row 1 versus the benchmark thresholds in Table 2, row 18). These findings support the possibility that dsEiger may signal through a heterotrimeric ligand architecture capable of engaging three identical dsGrnd receptors despite sequence divergence among the individual THDs.

**Figure 4:**
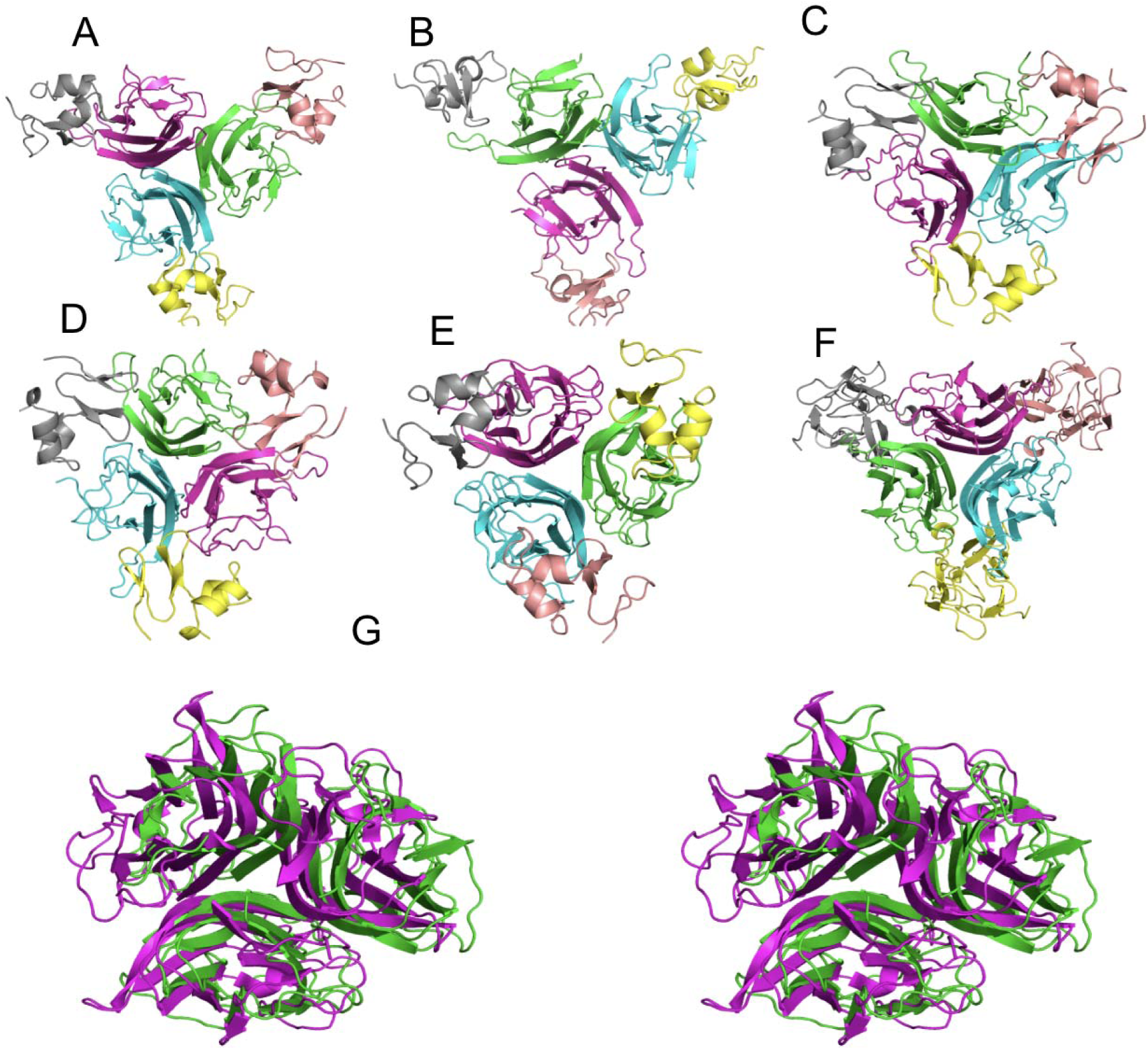
A-C) Predicted ligand-receptor complexes generated using the individual THDs (L1, L2, and L3) of dsEiger modeled as homotrimers in complex with dsGrnd respectively. D) Predicted ligand-receptor complex generated using all three THDs of dsEiger assembled as a heterotrimer in complex with dsGrnd. E) Crystal structure of the dmEiger-dmGrnd complex. F) Crystal structure of the human TNF-β receptor complex. G) Structural superposition of dmEiger homotrimer (green) with human TNF-α homotrimer (magenta), yielding an r.m.s.d. of 2.45 Å. Stereo representations corresponding to panels A-F are provided in Supplementary File 1, Figures S5a-f.

**Table 1:**
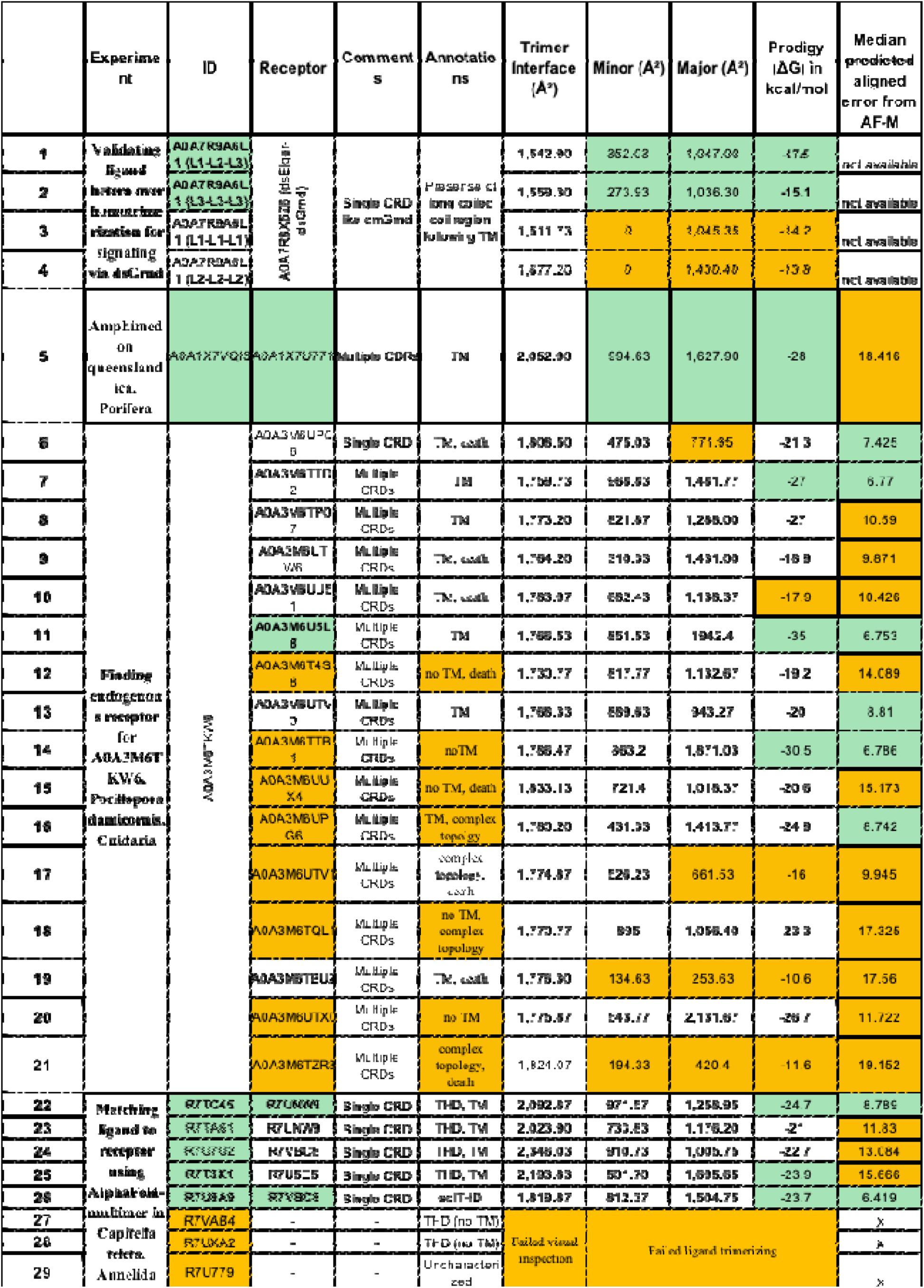
Parameters used for evaluation of predicted TNFSF-TNFRSF complexes, including interface area, binding affinity, binding configuration, and predicted aligned error (PAE). Columns 8 and 9 represent minor and major receptor-binding interfaces, respectively, based on the interaction geometry shown in Supplementary Figure S4e. Columns 10 and 11 represent ΔG and median PAE values obtained from AF-M predictions. Green indicates acceptable ranges, whereas orange indicates unacceptable ranges. The acceptable threshold for median PAE was set at <9.0.

**Table 2:**
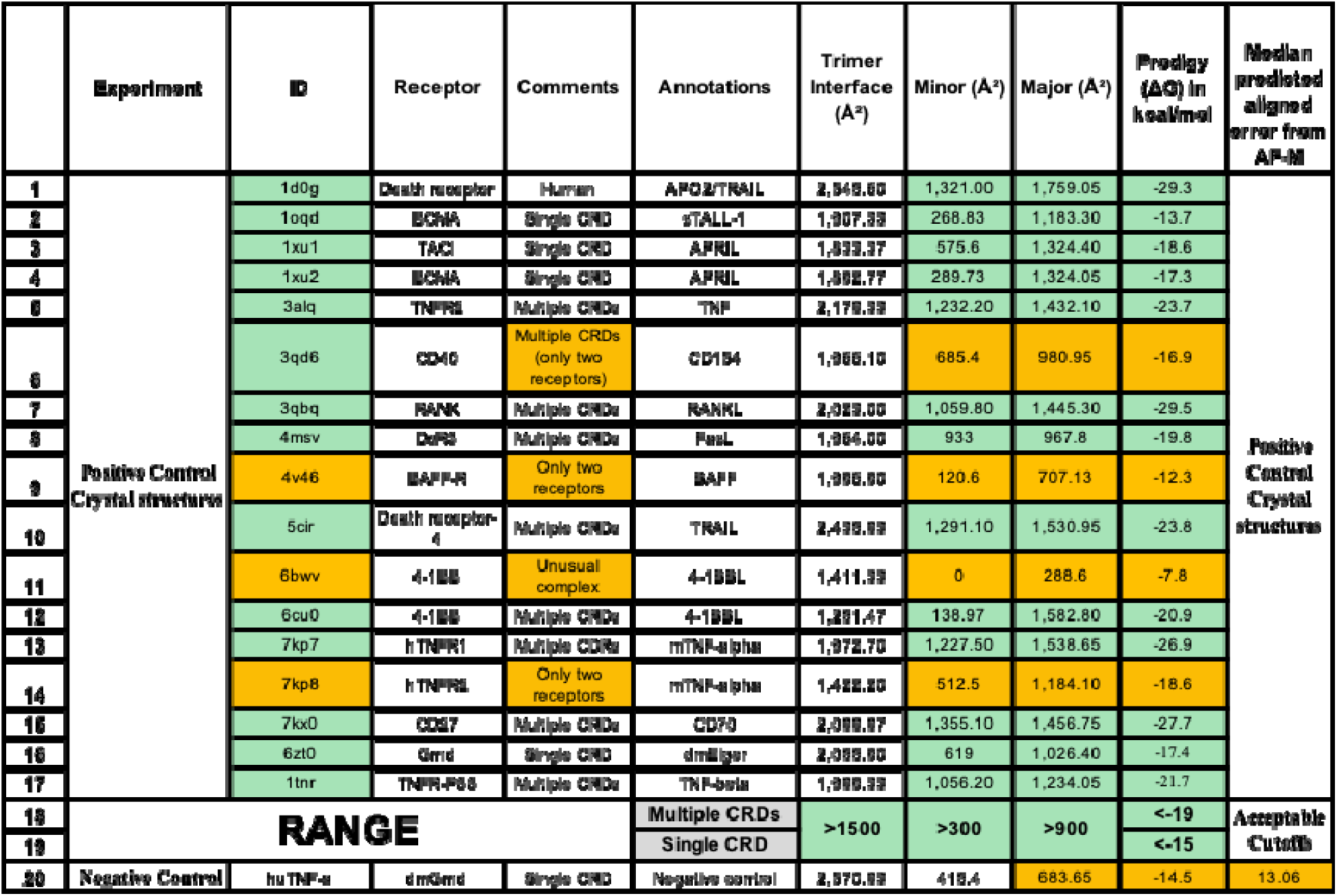
Parameters used for evaluation of predicted TNFSF-TNFRSF complexes, including interface area, binding affinity, binding configuration, and predicted aligned error (PAE). Columns 8 and 9 represent minor and major receptor-binding interfaces, respectively, based on the interaction geometry shown in Supplementary Figure S4e. Columns 10 and 11 represent ΔG and median PAE values obtained from AF-M predictions. Green indicates acceptable ranges, whereas orange indicates unacceptable ranges. The acceptable threshold for median PAE was set at <9.0.

In Figures 4a-c are models of ligand-receptor complexes formed by each of the three possible THDs of dsEiger as homotrimers interacting with dsGrnd. Among these, only the dsEiger-L3 homotrimer (corresponding to the C-terminal THD) displayed canonical receptor engagement, with dsGrnd binding at the trimeric ligand interface as shown in Figure 4c. This complex satisfied all structural parameters within the acceptable range (Table 1, row 2), except for a relatively stronger predicted binding affinity. In contrast, as summarized in Table 1 (rows 3-4), homotrimers formed by dsEiger-L1 and dsEiger-L2 failed to bind dsGrnd at the canonical ligand interface.

### Conservation of the signaling modality in lower-invertebrates

We modeled the only Eiger (aqEiger) and wengen (aqWgn) from *Amphimedon queenslandica* homologs identified in Porifera using AF-M. This complex satisfied all structural and energetic criteria derived from experimentally determined complexes (Table 1, row 5). The receptor bound symmetrically at the three ligand interfaces and displayed a binding affinity of -28 kcal/mol, near the lower end of the acceptable range. Coordinates for this complex are also provided in PDB format through GitHub.

In *Pocillopora damicornis* (Cnidaria), sequence-based analyses identified nine Eiger paralogs, although only one, A0A3M6TKW6 (pdEiger), satisfied the canonical architecture of an N-terminal TM followed by a single THD. In contrast, this species displayed a substantial expansion of wgn homologs, with 15 receptors identified, together with the earliest detected appearance of a single grnd homolog (pdGrnd) prior to the bilaterian split based on the current sampling of Porifera and Ctenophora.

To evaluate the ability of AF-M to assign cognate receptors, we modeled all 16 possible complexes between pdEiger and the available receptors. Analysis of the resulting complexes suggested that pdEiger may interact favorably with at least five receptors, with the strongest predicted interaction observed for A0A3M6U5L8 (pdWgn), which displayed a binding affinity of -35 kcal/mol (Table 1, row 11). Notably, this complex also exhibited the lowest median predicted aligned error (PAE) among all tested receptor combinations. The only grnd homolog, A0A3M6UP06 (pdGrnd), failed to satisfy all acceptance criteria despite containing a death domain (Table 1, row 6). Although the complex displayed a median acceptable PAE of 7.425 and a ΔG of -21 kcal/mol, the major interface area was only 771 Å² (column 9, row 6), suggesting that the threshold used for single-CRD receptors may be overly restrictive for definitive rejection.

Among the 15 wgn-like receptors, only nine possessed a C-terminal TM region, and four of these additionally contained intracellular death domains. Two of these receptors (Table 1, rows 7-8) satisfied most structural criteria and exhibited binding affinities of approximately -27 kcal/mol. In particular, A0A3M6TTD2 (Table 1, row 7) displayed a low median PAE of 6.7 and therefore remains a plausible endogenous receptor candidate for pdEiger. PAE heatmaps for all modeled complexes are provided in Supplementary Figure S7, and a comparison of ΔG versus PAE values is presented in Supplementary Table S5.

To investigate whether components of the TNF signaling pathway were already present prior to the bilaterian split, we analyzed representative species from the phyla Porifera and Cnidaria for homologs of genes associated with the TNF pathway. A total of 26 genes implicated in TNF signaling pathway were selected for analysis. Query sequences were derived from homologs identified in a malaria vect r strain exhibiting low vectorial capacity (ref).

For each query, sequence hits were filtered based on annotation matching with that of the query sequence, confirming the presence of homologs in species belonging to Porifera and Cnidaria (Table 2). Our analysis identified homologs for 25 out of the 26 TNF pathway genes within these early-diverging metazoan lineages in UniProt with 17 common to both Porifra and Cnidaria.

These findings suggest that a substantial portion of the TNF signaling machinery predates the bilaterian divergence and was likely already established in early metazoans. The presence of these conserved signaling components in basal animal phyla supports the hypothesis that TNF-mediated signaling originated early in animal evolution and later underwent functional expansion and diversification in vertebrates and invertebrates.

Table 2: The 26 pathway genes found in species (column 1) under Cnideria and Porifera as annotated in UniProt As shown in the pathway diagram, genes for many signaling routes are fully intact in both Porifera and Cnidaria suggesting mature signaling by Eiger genes as far back as in lower invertebrates.

### A choice of Grnd receptor as a norm under invertebrates

To further test the hypothesis that grnd-mediated signaling emerged early in invertebrates and later became predominant in Arthropoda, we modeled all possible combinations between eight Eiger paralogs and eight grnd paralogs from *Capitella teleta* (Annelida), generating 64 distinct complexes. This experiment was designed to evaluate whether AF-M could both authenticate Eiger homologs through stable homotrimer formation and assign potential endogenous receptors based on the structural parameters established in Table 1 (rows 22-29).

Five of the eight predicted Eiger homologs formed homotrimers with interface areas within the acceptable range. These ligands also bound receptors symmetrically at the ligand interfaces and displayed favorable binding affinities (Table 1, rows 22-26), with two complexes additionally showing acceptable median PAE values. Coordinates for all 64 modeled complexes, including those with non-canonical receptor configurations, are provided in PDB format (see Data Availability section). Collectively, these analyses enabled high-confidence assignment of cognate receptors for at least two Eiger homologs in *Capitella teleta* (Table 1, rows 22 and 26).

### A hypothesis towards genetically parsimonious solution triggering paralog/function expansion

In Figure 6b and 6c there are two examples of THDs that form heterodimers within the same gene, again missing the 2nd TM required to keep the two THDs floating extracellular. In these cases, the TMs are extending to form coiled coil, perhaps offering additional stability, as in the case of dsEiger (Figure 6a). According to TMHMM prediction, in the case of topologies shown in Figure 6b and 6c, there is a weak TM in the loop going into the cell after the THDs. We believe that, since the AlphaFold predicted structures used DeepTMHMM, which do not predict these regions as TM, the weak TMs are predicted as loops. We find an example (Figure 6d) where the entire complication of the TMs and loops passing back and forth through the membrane is mitigated by fusing the two THDs outside the membrane. These genes provide, perhaps, insights on the intermediate steps in the evolution towards highly genetically parsimonious homotrimerization of three copies of THDs from a single gene.

## DISCUSSION

The TNFSF-TNFRSF system has been extensively characterized in humans and other vertebrates because of its central role in immunity, apoptosis, inflammation, and therapeutic targeting. In this study, we analyzed near-complete genomes and proteomes from 148 species spanning all major invertebrate phyla, including lineages that diverged before the bilaterian split. This analysis led to the identification of 290 Eiger, 336 wengen, and 115 grnd homologs (Figure 1). Selected homologs were further investigated phylogenetically and structurally using AlphaFold-Multimer (AF-M) to explore the structure-function evolution of the TNFSF-TNFRSF signaling system across invertebrates. Homologs of both Eiger and grnd are absent in all sampled species belonging to Nematoda, Tardigrada, and the Chelicerata branch of Arthropoda, as well as in parasitic species under Platyhelminthes, as shown in Figure 1a and Supplementary Figure S6.

The Eiger and grnd phylogenies shown in Figures 2a and 2b exhibit clustering patterns that reflect both phylum-level and species-level organization, suggesting lineage-specific functional expansion of TNFSF and TNFRSF members. In addition, TNF homology domains (THDs) originating from the same gene frequently cluster together in the phylogeny, consistent with relatively recent in-frame duplication events. For example, within Clade 2, several Eiger homologs from *Lottia gigantea* and *Octopus vulgaris* cluster together. Similarly, all TNFSF homologs from *Macrostomum lignano*, a member of Platyhelminthes, form a well-supported cluster within Clade 7 (84% confidence). At the domain level, the two THD domains from A0A8B8AD19 in *Crassostrea virginica* cluster closely with THDs from V4AUT9 and V4BF89 in *Lottia gigantea* with 100% confidence, further supporting recent domain duplication events within these lineages.

Among lower invertebrates, no homologs of Eiger or grnd were detected in the sampled species of Ctenophora, the proposed sister group to Metazoa (Schultz et al. 2023). The earliest appearance of grnd homologs in our dataset was observed in Cnidaria. Based on the current sampling, within Porifera, we identified homologs of Eiger and wgn but not grnd. Importantly, structural modeling of the only identified ligand-receptor pair in *Amphimedon queenslandica* (Porifera) (aqEiger and aqWgn) produced a highly favorable complex (Table 1, row 5), supporting the existence of a potentially functional TNFSF-TNFRSF configuration in Porifera.

A particularly notable pattern emerged in *Pocillopora damicornis* (Cnidaria), where we observed a substantial expansion of wgn homologs together with the presence of a single grnd homolog. Table 1 (rows 26-41) summarizes the structural and energetic parameters for complexes formed between the sole Eiger homolog, pdEiger, complexed with all identified receptor homologs. While the complex of pdEiger and pdGrnd (Table 1, row 6) was not too week for a definitive rejection. In contrast, several wgn homologs interacted favorably with pdEiger, with A0A3M6U5L8 (pdWgn) emerging as the strongest candidate cognate receptor (Table 1, row 11).

Taken together, AF-M-based analyses in lower invertebrates support the hypothesis that ancestral TNFSF signaling was primarily mediated through wgn-like receptors, extending at least as far back as Porifera based on the current sampling. These observations further suggest that grnd-based signaling emerged later during invertebrate evolution and subsequently became dominant within Arthropoda.

An especially intriguing case was observed in *Darwinula stevensoni*, an Arthropoda, where the only identified TNFSF homolog, dsEiger, contains three tandem THD domains within a single gene, while the genome encodes only one dsGrnd receptor. This architecture suggests the possibility of signaling through heterotrimerization of the three distinct THDs encoded within dsEiger. Using AF-M together with the structural filtering criteria established in Table 2 (rows 18-19), we demonstrate that dsEiger is capable of engaging dsGrnd both through heterotrimerization involving the three distinct THDs (L1, L2, and L3) and through homotrimerization involving the terminal THD-L3. These observations support the possibility that heterotrimeric TNFSF signaling may represent an ancient or transitional signaling strategy retained in specific invertebrate lineages.

In *Capitella teleta* (Annelida), which contains multiple Eiger homologs together with a substantial expansion of grnd paralogs, we identified a useful model for studying the emergence and stabilization of grnd-mediated signaling following the bilaterian split. Using AF-M, we modeled all sixty-four possible ligand-receptor combinations involving THD domains from eight Eiger homologs and CRD domains from eight grnd homologs. The five authentic Eiger homologs could be assigned endogenous receptors. Notably, several receptors interacted favorably with more than one ligand, a feature that is not uncommon among vertebrate TNFSF-TNFRSF systems.

Function expansion by in-frame duplication of Eiger and receptor paralogs within a single gene presents a significant topological challenge because members of TNFSF and TNFRSF typically possess obligate N-terminal or C-terminal transmembrane regions. Such arrangements would place alternating extra-cellular binding domains on opposite sides of the membrane, imposing severe structural and evolutionary constraints during functional expansion (Duran and Meiler, 2013). One striking example is illustrated in Figure 4b, where the three repeating arrangements of transmembrane regions (red), THDs (triangles), and linker loops (lines) in dsEiger lack the additional transmembrane segments theoretically required to orient each THD extracellularly (as depicted schematically in Figure 5a). Instead, the three TM regions are extended by coiled-coil domains projecting into the extracellular matrix, potentially stabilizing the TM helices in a parallel bundle configuration and thereby suspending the heterotrimeric ligand extracellularly for signaling.

**Figure 5:**
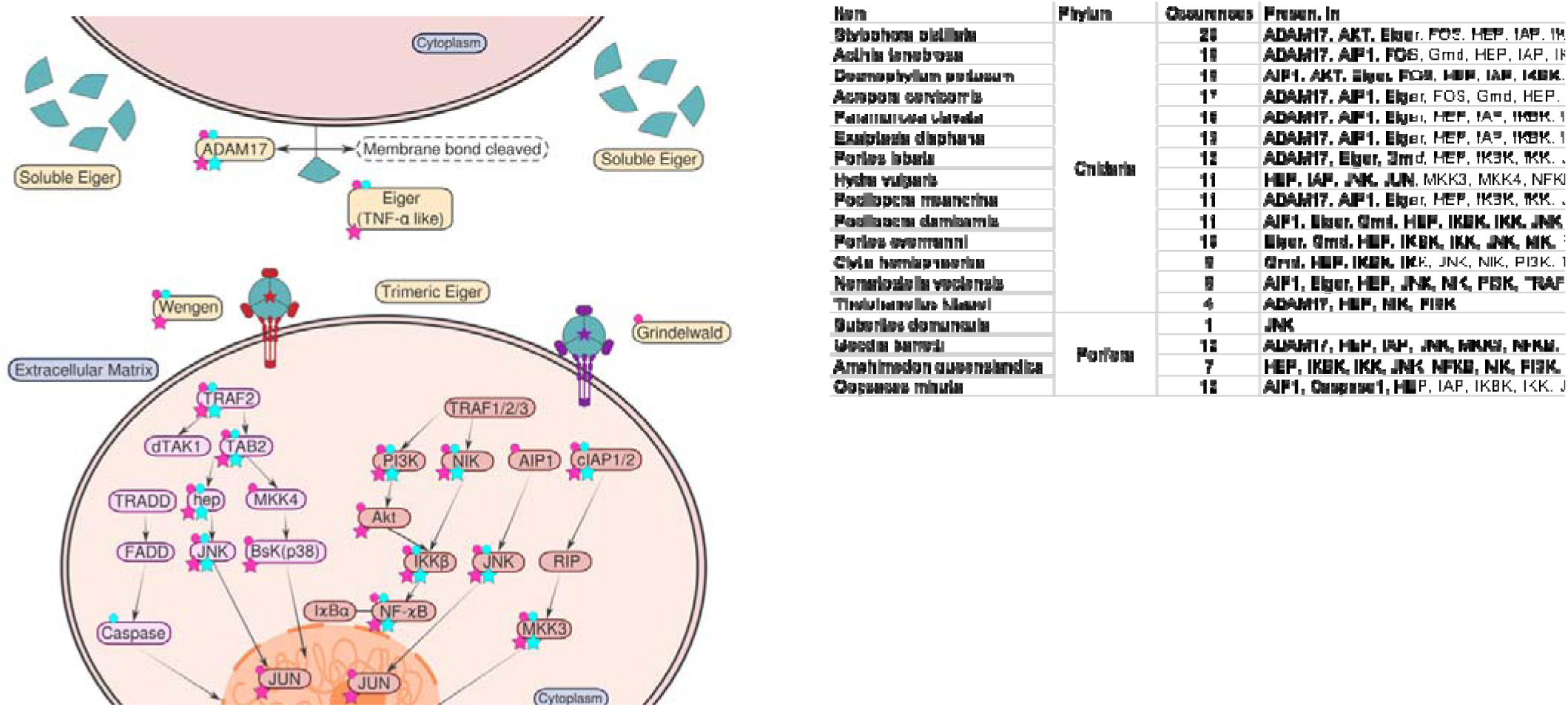
Homologs of the 26 TNF signaling pathway genes highlighted in Porifera (cyan) and Cnidaria (magenta).

**Figure 6:**
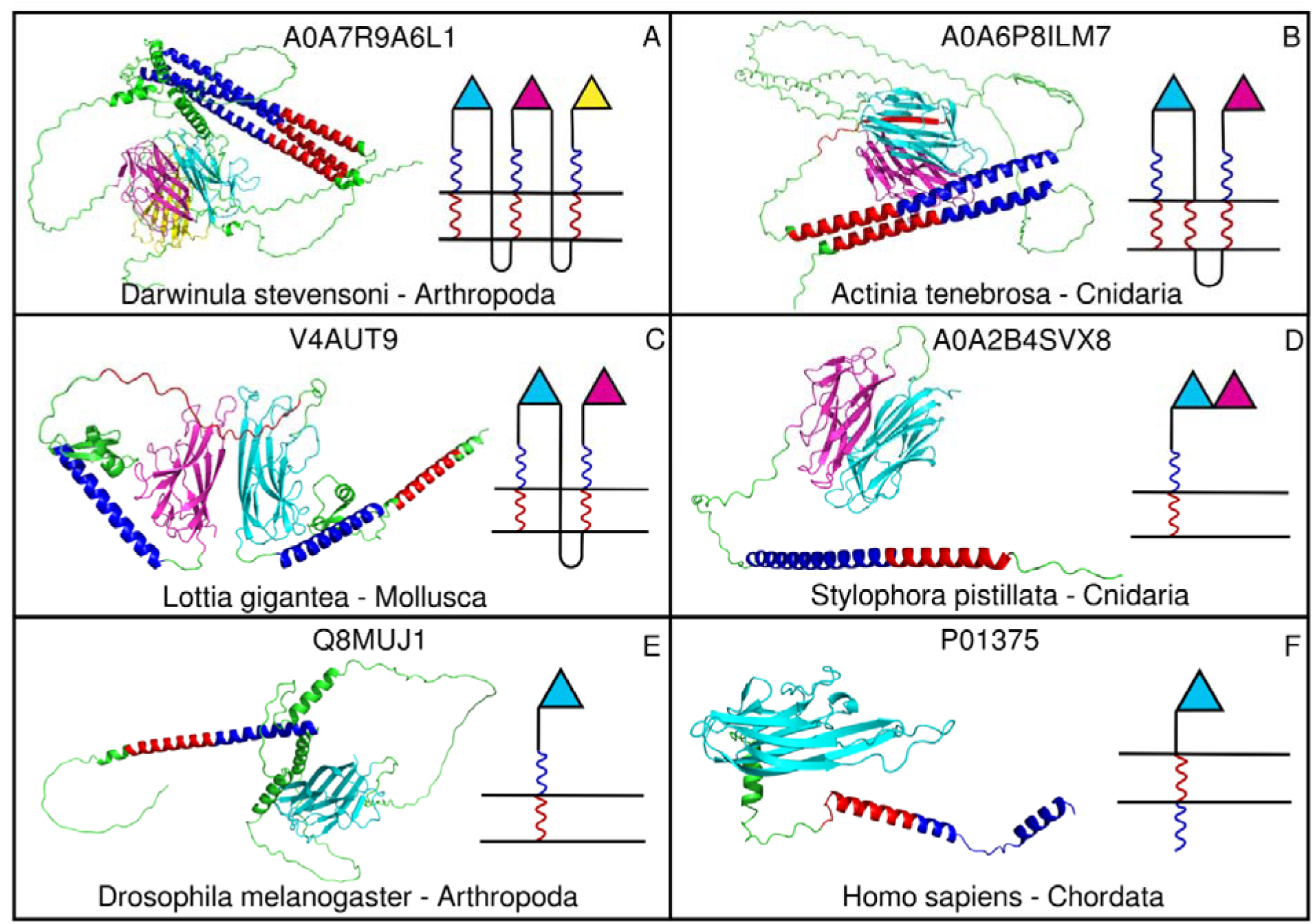
A-D) 3D topologies of rare surviving homologs of TNFSF containing multiple Eiger-like domains (pink, blue and green) within single genes under Cnidaria, Mollusca, and Arthropoda. **E-F)** topologies of the prevalent protein architecture in D. melanogaster and human paralogs. THDs are in triangle, transmembrane regions are in red and coiled coil in blue.

The complex topologies of TNFSF homologs identified across diverse invertebrate lineages allowed us to reconstruct potential evolutionary intermediates that may represent successive stages in the emergence and diversification of TNF signaling systems. In particular, the presence of N-terminal transmembrane (TM) domains in several homologs suggests a structural constraint during early evolution, as such configurations could invert the orientation of extracellular domains into the intracellular space, thereby complicating functional expansion through simple gene duplication alone. Based on these observations, we hypothesize that the intermediate topologies retained in extant species reflect evolutionary transitions that ultimately led to a genetically parsimonious solution for both ligand and receptor diversification. This solution may have involved the emergence of homo-oligomerization strategies that enabled functional diversification without extensive structural redesign of ligand and receptor architectures. Furthermore, the utilization of Grindelwald (grnd)-like receptors containing a single cysteine-rich domain (CRD) prior to the major radiation of Arthropoda suggests that simplified receptor architectures may have preceded later expansions and specialization events. Collectively, these findings support a model in which TNFSF and TNFRSF evolution proceeded through structurally constrained intermediates before undergoing extensive functional expansion in vertebrates, including humans.

## MATERIALS AND METHODS

### Materials

Reference proteomes were selected from the <u>UniProt database</u> to represent a broad cross-section of taxonomic diversity within UniProtKB. Proteomes with BUSCO scores above 90% were preferentially selected as indicators of near-complete proteome coverage. In phyla such as Platyhelminthes and Tardigrada, where only a limited number of reference proteomes were available, additional proteomes with BUSCO scores above 85% were included to improve representation. For Porifera, where only a single reference proteome of *Amphimedon queenslandica* was available in UniProt, a second proteome from Ephydatia muelleri was obtained from <u>EphyBase</u>. On the other hand, in phyla containing many completed genomes, representative species from distinct genera were selected to maximize phylogenetic diversity while avoiding redundancy. For example, within Nematoda, only 11 of the 128 species with completed genomes were included without compromising taxonomic breadth.

### Building HMM models for invertebrates

A major challenge in identifying homologs and paralogs of TNFSF and TNFRSF families is their extensive sequence divergence across evolutionary lineages. Conventional sequence similarity approaches such as BLAST produce substantial numbers of false positives and false negatives for these families. To improve sensitivity and specificity, we initially used existing HMM profiles for TNFSF (PF00229.21) and TNFRSF (PF00020.22) to search UniProt proteomes. However, based on known members characterized in the literature, these models showed insufficient sensitivity for detecting invertebrate homologs. For instance, the TNFRSF HMM profile (PF00020.21) failed to detect known grnd homologs in both *Drosophila* and malaria vectors, the only invertebrates in which TNFRSF members have been experimentally characterized.

To overcome these limitations, we iteratively refined and rebuilt HMM models optimized for vertebrate homolog detection by adding known invertebrate homolods to the seed alignment from Pfam HMM profiles for TNFSF (PF00229.21) and TNFRSF (PF00020.22) using MUSCLE implemented in MEGA. Conserved regions were manually inspected to remove poorly aligned segments and excessive gaps. The curated multiple sequence alignments (MSAs) were then used to build invertebrate-specific HMMs using `hmmbuild` from HMMER v3.3.

As more homologs were found they were added to the seed alignment iteratively. This iterative procedure was repeated until no additional bona fide homologs were identified across the sampled species. Following the success of the in-house grnd HMM, a similar strategy was adopted to build improved HMMs for Eiger homolog detection to minimize false negatives caused by extensive sequence divergence among invertebrates.

The resulting in-house models achieved negligible false positive and false negative rates, substantially increasing confidence in the mining pipeline (Supplementary Figure S2; see Data Availability section). Additional structural filters were applied to eliminate unrelated proteins sharing superficial domain similarity. For example, authentic TNFSF homologs were required to possess a transmembrane region located approximately 50-100 amino acids N-terminal to the folding domain. Similarly, putative TNFRSF homologs were filtered by requiring a transmembrane segment positioned within 80-120 amino acids of the cysteine-rich domains (CRDs), thereby excluding unrelated cysteine-rich proteins.

### Evaluation of in-house built HMM models

The performance of PFam-derived and in-house HMM models was evaluated statistically (Supplementary Figure S2). Metrics included sensitivity, specificity, false-positive rate, false-negative rate, predictive values, and likelihood ratios. The custom invertebrate HMMs are available through the GitHub repository listed in the Data Availability section.

The following formulas were used:

- Sensitivity = TNF/TNFR positive hits / total TNF/TNFR hits
- Specificity = non-TNF/TNFR negative hits / total non-TNF/TNFR hits
- False positives = non-TNF/TNFR positive hits / total non-TNF/TNFR hits
- False negatives = TNF/TNFR positive hits / total TNF/TNFR hits
- Positive predictive value = true positive hits / total positive hits
- Negative predictive value = true negative hits / total negative hits
- Positive likelihood ratio = sensitivity / (1 − specificity)
- Negative likelihood ratio = (1 − sensitivity) / specificity

### Authentication of Presence/Absence Calls

As emphasized in evolutionary analyses of signaling systems, accurate presence/absence assignment is critical for reconstructing evolutionary history. To reduce false-positive assignments, putative TNFSF and TNFRSF homologs were further filtered using topology constraints characteristic of type-II and type-I transmembrane proteins, respectively.

Members of the TNFSF family display a characteristic topology consisting of an N-terminal transmembrane (TM) region and an extracellular C-terminal TNF Homology Domain (THD). The THD contains conserved sequence signatures, including GLY and FFG motifs, that are associated with oligomerization and receptor binding during signaling.

TNFRSF homologs, including TNFR1- and TNFR2-like receptors, exhibit an extracellular domain composed of 1-6 cysteine-rich domains (CRDs). These CRDs are stabilized by disulfide bridges surrounding a conserved core motif. Typically, only one CRD directly participates in ligand binding and commonly carries a CXXCXXC or CXXCXXXC signature. Many TNFRSF homologs also contain an intracellular death domain (DD), which mediates downstream signaling.

Predicted TM regions were identified using DeepTMHMM v1.0.24 and TMHMM. Structural validation was performed using AlphaFold-predicted structures available through UniProt. Where AlphaFold models were unavailable, structures were generated using the <u>RoseTTAFold server</u>.

Predicted THD and CRD structures were structurally superimposed onto experimentally solved complexes, including PDB:1tnr and PDB:6zt0. Authentication was based on low RMSD and preservation of characteristic fold topology.

To further improve sensitivity and validate absence calls, Foldseek v8 searches were performed against the AlphaFold/UniProt structural database containing more than 214 million predicted protein structures. The 3D structures of dmEiger, dmGrnd, and dmWgn domains from Drosophila melanogaster were used as queries. Foldseek searches were performed in TMalign mode with a maximum of 5000 hits and phylum-specific taxonomic filters. Hits were filtered based on taxonomy, structural coverage, RMSD (<4 Å), transmembrane topology, and functional relevance. Entries from Bacteria, Archaea, and non-metazoan groups were manually inspected to eliminate structurally similar but functionally unrelated proteins.

### Phylogenetic Analysis and Reconciliation

The THD and grnd homology domains from all bona fide homologs were extracted and combined with homologous domains from 19 human TNFSF paralogs and 9 human TNFRSF members containing grnd-like homology domains.

Multiple sequence alignments were generated using MAFFT v7.520. Maximum-likelihood phylogenetic trees were constructed using IQ-TREE v1.6.12. The optimal amino acid substitution model was selected using ModelFinder, and branch support was estimated using ultrafast bootstrap analysis with 1,000 replicates.

### Evaluating AF-M for Modeling TNFSF-TNFRSF Complexes

In CASP16, AlphaFold-Multimer (AF-M) performed only marginally better for de novo prediction of protein complexes (Zhang et al. 2025). However, the TNFSF-TNFRSF system represents a fundamentally different benchmark because more than 17 experimentally determined ligand-receptor complexes are available in the Protein Data Bank (PDB). In deep-learning approaches, predictive accuracy improves substantially with the availability of related template structures; therefore, AF-M performance is expected to vary considerably across protein systems. Our previous work demonstrated that AlphaFold functions largely as a large-scale homology modeling framework (Goswami and Srinivasan 2025). Preliminary analyses from our laboratory further suggest that AF-M leverages homology in higher-order template configurations before refinement through docking. Here, we exploited failures to preserve canonical ligand-receptor configurations as a criterion for visually filtering non-canonical complexes.

To establish criteria for accepting or rejecting predicted TNFSF-TNFRSF complexes, we derived a set of structural and energetic parameters from experimentally determined 3:3 ligand-receptor complexes involving 15 human TNFSF paralogs and dmEiger bound to their cognate receptors. These parameters included:

- homotrimerization interface area,
- ligand-receptor interface area,
- mode and symmetry of receptor binding,
- receptor folding and disulfide topology,
- threefold symmetry of receptor engagement, and
- binding affinity.

Rows 1-17 in Table 2 summarize these parameters for experimentally validated complexes, while rows 18-19 define acceptable thresholds for evaluating AF-M predictions. Based on these benchmarks, acceptable complexes were required to exhibit:

- ligand homotrimer interfaces >1500 Å²,
- minor and major receptor-binding epitopes >300 Å² and >900 Å², respectively, and
- binding affinities stronger than approximately -15 to -19 kcal/mol.

A limitation in deriving acceptable ranges for binding affinity and median predicted aligned error (PAE) values in Table 2 is that these thresholds do not account for differences in physiological state, receptor competition, and signaling complexity between simple and highly complex organisms. In addition, the PAE values reported by AlphaFold-Multimer (AF-M) represent residue-pair confidence scores on a scale ranging from 0-30. For the purposes of this study, a median PAE threshold of <9.0 was considered acceptable for predicted complexes.

Custom Python scripts utilizing Biopython were developed to quantify ligand-ligand and ligand-receptor interaction interfaces from PDB files. Atomic contacts between chains were identified using a KDTree algorithm with a 3.5 Å distance cutoff. Interactions were catalogued for all inter-chain combinations among TNFSF trimer chains (A, B, C) and TNFRSF receptor chains (D, E, F). Residue-level interactions were subsequently consolidated into grouped interaction summaries. Binding affinities (ΔG; kcal mol^-1^) for ligand-receptor complexes were calculated using the <u>PRODIGY</u> <u>server</u>.

To evaluate the usefulness of these parameters, we modeled a negative-control complex between human TNF-α and dmGrnd from fly using AF-M. Although the three huTNF-α chains formed favorable receptor-binding epitopes, the receptors bound in asymmetric and non-canonical orientations. The computed parameters for this complex (Table 2, row 20) also failed to satisfy the acceptance criteria listed in row 19. Coordinates of the modeled structure are available through GitHub (see Data Availability section).

### AlphaFold and AlphaFold-Multimer

Structure predictions were generated using LocalColabFold v1.5.5, compatible with AlphaFold v2.3.2. ColabFold uses MMseqs2 for homology detection against ColabFoldDB, which integrates UniRef30, BFD, MGnify, and related databases.

AlphaFold-Multimer (AF-M) was used to model ligand-receptor complexes. Complex evaluation incorporated:

- ligand homotrimer interface area
- receptor binding interface area
- binding affinity (ΔG)
- receptor orientation and symmetry
- disulfide topology
- predicted aligned error (PAE)

Experimentally determined TNFSF-TNFRSF complexes from vertebrates and the dmEiger-dmGrnd complex served as calibration standards for acceptable parameter ranges.

## ACKNOWLEDGEMENTS

The authors acknowledge the Government of Karnataka for supporting the computational infrastructure at the Institute of Bioinformatics and Applied Biotechnology and for funding MG and MKG through the BioIT initiative. The authors also acknowledge support from DBT Builder Sanction No. dated 08/03/2022 and CCB Sanction No. BT/PR40212/BTIS/137/40/2022 dated 19/12/2022.

## AUTHOR CONTRIBUTION

Kaushiki: Tracking TNF-TNFR system under invertebrate and before the split; MG: Tracking TNF- TNFR system under other branches of life and calculating parameters to evaluate 3D complexes built using AlphaFold and help with the manuscript; NM: Tracking TNF-TNFR system under Arthropoda including identification of Eiger and grnd in *Darwinula stevensoni*;; AS: Building/evaluating models of complexes using AF-M; MKG: Generation and evaluation of 140 complexes using AlphaFold-multimer and aiding with Figures; SS: Conceptualization, execution and writing of the manuscript.

## DATA AVALABILITY

- The high-resolution phylogenetic tree of 191 Eiger homologs from invertebrates and corresponding grnd genes are available in PDF format under Supplementary Files S2 and S3.
- HMM models for TNF/Eiger and grnd built in-house can be downloaded from https://github.com/moushmigoswami/TNF/tree/main/Supplementary/invertebrates_HMM_models.
- Highly annotated phyla-wise table homologs/paralogs from species identified by mining 148 completed genomes under both invertebrate and lower invertebrate in excel format can be found https://github.com/moushmigoswami/TNF/blob/main/Supplementary/Table_Eiger_Wgn_Grnd_Homology.xlsx.
- The 3D structures of all the 140 complexes including those listed in Table 2 in PDB format and those used to screen for inhibitors against huTNF-α using the 96 CRD doans from grnd gene from Arthropoda can be found https://github.com/moushmigoswami/TNF/tree/main/3D_PDB_TNF_TNFR_Complexes. The PDB structure of complexes can be viewed using PyMOL or Chimera (Pettersen et al. 2004).
- A domain-based color-coded PyMOL Session Files for figures in Supplementary Figure S4 can be found https://github.com/moushmigoswami/TNF/tree/main/Supplementary/S4_Fig_PSE_Files.
- Programs to compute binding interfaces to test reproducibility. https://github.com/moushmigoswami/TNF/tree/main/Supplementary/Interface_Area_Calculation/Intra_Ligand_Interface https://github.com/moushmigoswami/TNF/tree/main/Supplementary/Interface_Area_Calculation/Liga nd_Receptor_Interface
- MSA for generating HMM profile:

https://github.com/moushmigoswami/TNF/tree/main/Supplementary/MSA

## BIBLIOGRAPHY

1. Barrio-Hernandez I, Yeo J, Jänes J, et al (2023) Clustering predicted structures at the scale of the known protein universe. Nature 622:637–645. 10.1038/s41586-023-06510-w

2. Barshis DJ, Ladner JT, Oliver TA, et al (2013) Genomic basis for coral resilience to climate change. Proc Natl Acad Sci U S A 110:1387–1392. 10.1073/pnas.1210224110

3. Bertevello CR, Russo BRA, Tahira AC, et al (2020) The evolution of TNF signaling in platyhelminths suggests the cooptation of TNF receptor in the host-parasite interplay. Parasit Vectors 13:491. 10.1186/s13071-020-04370-1

4. Bertinelli M, Paesen GC, Grimes JM, Renner M (2019) High-resolution crystal structure of arthropod Eiger TNF suggests a mode of receptor engagement and altered surface charge within endosomes. Commun Biol 2:293. 10.1038/s42003-019-0541-0

5. Bodmer J-L, Schneider P, Tschopp J (2002) The molecular architecture of the TNF superfamily. Trends Biochem Sci 27:19–26. 10.1016/s0968-0004(01)01995-8

6. Cock PJA, Antao T, Chang JT, et al (2009) Biopython: freely available Python tools for computational molecular biology and bioinformatics. Bioinformatics 25:1422–1423. 10.1093/bioinformatics/btp163

7. Collette Y, Gilles A, Pontarotti P, Olive D (2003) A co-evolution perspective of the TNFSF and TNFRSF families in the immune system. Trends Immunol 24:387–394. 10.1016/s1471-4906(03)00166-2

8. De Zoysa M, Jung S, Lee J (2009) First molluscan TNF-alpha homologue of the TNF superfamily in disk abalone: molecular characterization and expression analysis. Fish Shellfish Immunol 26:625–631. 10.1016/j.fsi.2008.10.004

9. Duran AM, Meiler J (2013) Inverted topologies in membrane proteins: a mini-review. Comput Struct Biotechnol J 8:e201308004. 10.5936/csbj.201308004

10. Eddy SR (1998) Profile hidden Markov models. Bioinformatics 14:755–763. 10.1093/bioinformatics/14.9.755

11. Goswami M, Srinivasan S (2025) Tracing the function expansion for a primordial protein fold in the era of fold-based function prediction: beta-trefoil. 2025.02.18.638835

12. Hoang DT, Chernomor O, von Haeseler A, et al (2018) UFBoot2: Improving the Ultrafast Bootstrap Approximation. Mol Biol Evol 35:518–522. 10.1093/molbev/msx281

13. Igaki T, Kanda H, Yamamoto-Goto Y, et al (2002) Eiger, a TNF superfamily ligand that triggers the Drosophila JNK pathway. EMBO J 21:3009–3018. 10.1093/emboj/cdf306

14. Jumper J, Evans R, Pritzel A, et al (2021) Highly accurate protein structure prediction with AlphaFold. Nature 596:583–589. 10.1038/s41586-021-03819-2

15. Kalyaanamoorthy S, Minh BQ, Wong TKF, et al (2017) ModelFinder: fast model selection for accurate phylogenetic estimates. Nat Methods 14:587–589. 10.1038/nmeth.4285

16. Kanda H, Igaki T, Kanuka H, et al (2002) Wengen, a member of the Drosophila tumor necrosis factor receptor superfamily, is required for Eiger signaling. J Biol Chem 277:28372–28375. 10.1074/jbc.C200324200

17. Kim DE, Chivian D, Baker D (2004) Protein structure prediction and analysis using the Robetta server. Nucleic Acids Res 32:W526–531. 10.1093/nar/gkh468

18. Krupovic M, Koonin EV (2017) Multiple origins of viral capsid proteins from cellular ancestors. Proc Natl Acad Sci U S A 114:E2401–E2410. 10.1073/pnas.1621061114

19. Li L, Qiu L, Song L, et al (2009) First molluscan TNFR homologue in Zhikong scallop: molecular characterization and expression analysis. Fish Shellfish Immunol 27:625–632. 10.1016/j.fsi.2009.07.009

20. Majoros WH, Pertea M, Delcher AL, Salzberg SL (2005) Efficient decoding algorithms for generalized hidden Markov model gene finders. BMC Bioinformatics 6:16. 10.1186/1471-2105-6-16

21. Marín I (2020) Tumor Necrosis Factor Superfamily: Ancestral Functions and Remodeling in Early Vertebrate Evolution. Genome Biol Evol 12:2074–2092. 10.1093/gbe/evaa140

22. Marín I (2025) Vertebrate TNF Superfamily: Evolution and Functional Insights. Biology (Basel) 14:54. 10.3390/biology14010054

23. Mirdita M, Schütze K, Moriwaki Y, et al (2022) ColabFold: making protein folding accessible to all. Nat Methods 19:679–682. 10.1038/s41592-022-01488-1

24. Mitternacht S (2016) FreeSASA: An open source C library for solvent accessible surface area calculations. F1000Res 5:189. 10.12688/f1000research.7931.1

25. Nguyen L-T, Schmidt HA, von Haeseler A, Minh BQ (2015) IQ-TREE: a fast and effective stochastic algorithm for estimating maximum-likelihood phylogenies. Mol Biol Evol 32:268–274. 10.1093/molbev/msu300

26. Palmerini V, Monzani S, Laurichesse Q, et al (2021) Drosophila TNFRs Grindelwald and Wengen bind Eiger with different affinities and promote distinct cellular functions. Nat Commun 12:2070. 10.1038/s41467-021-22080-9

27. Pettersen EF, Goddard TD, Huang CC, et al (2004) UCSF Chimera--a visualization system for exploratory research and analysis. J Comput Chem 25:1605–1612. 10.1002/jcc.20084

28. Pozzolini M, Scarfì S, Ghignone S, et al (2016) Molecular characterization and expression analysis of the first Porifera tumor necrosis factor superfamily member and of its putative receptor in the marine sponge Chondrosia reniformis. Dev Comp Immunol 57:88–98. 10.1016/j.dci.2015.12.011

29. Quistad SD, Stotland A, Barott KL, et al (2014) Evolution of TNF-induced apoptosis reveals 550 My of functional conservation. Proc Natl Acad Sci U S A 111:9567–9572. 10.1073/pnas.1405912111

30. Quistad SD, Traylor-Knowles N (2016) Precambrian origins of the TNFR superfamily. Cell Death Discov 2:16058. 10.1038/cddiscovery.2016.58

31. Schultz DT, Haddock SHD, Bredeson JV, et al (2023) Ancient gene linkages support ctenophores as sister to other animals. Nature 618:110–117. 10.1038/s41586-023-05936-6

32. Simão FA, Waterhouse RM, Ioannidis P, et al (2015) BUSCO: assessing genome assembly and annotation completeness with single-copy orthologs. Bioinformatics 31:3210–3212. 10.1093/bioinformatics/btv351

33. Sonar S, Lal G (2015) Role of Tumor Necrosis Factor Superfamily in Neuroinflammation and Autoimmunity. Front Immunol 6:364. 10.3389/fimmu.2015.00364

34. Srinivasan S, Ghosh C, Das S, et al (2022) Identification of a TNF-TNFR-like system in malaria vectors (Anopheles stephensi) likely to influence Plasmodium resistance. Sci Rep 12:19079. 10.1038/s41598-022-23780-y

35. Steichele M, Sauermann LS, König A-C, et al (2021) Ancestral role of TNF-R pathway in cell differentiation in the basal metazoan Hydra. J Cell Sci 134:jcs255422. 10.1242/jcs.255422

36. Tian T, Dubin K, Jin Q, et al (2012) Disruption of TNF-α/TNFR1 function in resident skin cells impairs host immune response against cutaneous vaccinia virus infection. J Invest Dermatol 132:1425–1434. 10.1038/jid.2011.489

37. Tuliuan J, Javier W, Austria E, Valentino M (2025) Evolutionary relationship of edible mollusks based on phylogenetic analysis using 16s RRNA and COX1 gene sequences. Journal of Applied Biological Sciences 19:97–104. 10.71336/jabs.1456

38. van Kempen M, Kim SS, Tumescheit C, et al (2024) Fast and accurate protein structure search with Foldseek. Nat Biotechnol 42:243–246. 10.1038/s41587-023-01773-0

39. Wu Y, He J, Yao G, et al (2020) Molecular cloning, characterization, and expression of two TNFRs from the pearl oyster Pinctada fucata martensii. Fish Shellfish Immunol 98:147–159. 10.1016/j.fsi.2020.01.010

40. Xue LC, Rodrigues JP, Kastritis PL, et al (2016) PRODIGY: a web server for predicting the binding affinity of protein-protein complexes. Bioinformatics 32:3676–3678. 10.1093/bioinformatics/btw514

41. Zhang J, Yuan R, Kryshtafovych A, et al (2025) Assessment of Protein Complex Predictions in CASP16: Are we making progress? bioRxiv 2025.05.29.656875. 10.1101/2025.05.29.656875

42. Zhang L, Li L, Guo X, et al (2015) Massive expansion and functional divergence of innate immune genes in a protostome. Sci Rep 5:8693. 10.1038/srep08693

