## Supplementary Material for "TNF-TNFR Signaling Modality Conserved Across Invertebrate"

**Supplementary Table S1: Summary of proteome analysis for invertebrate Phylum**

Column 1 shows the phyla that have been studied, column 2 shows the total number of proteomes available in the UniProt database, column 3 shows the number of reviewed proteomes available and column 5 shows the number of proteomes that have been taken for this study. Proteomes of different species from various phyla having a BUSCO completeness of 85% or higher have been studied.

| **Phylum** | **Total proteomes** | **Reviewed proteome in UniProt** | **Proteomes with BUSCO C >= 85%** | **Proteomes studied** |
| --- | --- | --- | --- | --- |
| Tardigrada | 2 | 2 | 1 | 2 (one non-reference proteome with the highest BUSCO) |
| Nematoda | 128 | 62 | 16 | 11 (only single proteome with the highest BUSCO under each genus) |
| Mollusca | 17 | 11 | 4 | 4 |
| Annelida | 6 | 3 | 3 | 3 |
| Brachiopoda | 1 | 1 | 1 | 1 |
| Platyhelminthes | 49 | 28 | 1 | 2 (one non-reference proteome with the highest BUSCO) |
| Cnidaria | 10 | 7 | 5 | 5 |
| Porifera | 2 | 1 | 1 | 2 (1 from EphyBase) |
| Arthropoda | 246 | 169 | 116 | 118 ( two Proteomes with a lower BUSCO) |
| **Total** | **542** | **355** | **157** | **157** |

**Supplementary Table S2: Summary of the number of Eiger, Wgn and Grd homologs per species per phylum**

Column 1 shows the name of the species from each phylum. Columns 2, 3 and 4 show the number of Eiger, Wgn and Grnd homologs in each species respectively.

| **Phyla** | **Eiger** | | **Wgn** | | **Grnd** | |
| --- | --- | --- | --- | --- | --- | --- |
|  | **# of hits using in-house HMM** | **# of hits after filtering** | **# of hits using in-house HMM** | **# of hits after filtering** | **# of hits using in-house HMM** | **# of hits after filtering** |
| **Tardigrada** | 3 | 0 | 5 | 4 | 3 | 0 |
| **Nematoda** | 18 | 0 | 26 | 5 | 7 | 0 |
| **Mollusca** | 216 | 63 | 92 | 44 | 46 | 4 |
| **Annelida** | 56 | 22 | 28 | 5 | 24 | 14 |
| **Brachiopoda** | 41 | 17 | 52 | 11 | 12 | 6 |
| **Platyhelminthes** | 22 | 6 | 44 | 19 | 1 | 1 |
| **Cnidaria** | 71 | 32 | 154 | 90 | 34 | 2 |
| **Porifera** | 6 | 2 | 16 | 3 | 2 | 0 |
| **Arthropoda** | 148 | 148 | 155 | 155 | 88 | 88 |
| **Total** | **581** | **290** | **572** | **336** | **217** | **115** |

**Supplementary Table S3: Table containing the number of initial hits from initial HMM search and those considered positive hits after multiple biological/structural filters.**

| **PHYLUM PORIFERA** | | | |
| --- | --- | --- | --- |
| **Species** | **Eiger** | **Wgn** | **Grnd** |
| *Amphimedon queenslandica* | 1 | 2 | 0 |
| *Ephydatia murlleri* | 1 | 1 | 0 |
| **Total** | **2** | **3** | **0** |

| **PHYLUM CNIDARIA** | | | |
| --- | --- | --- | --- |
| **Species** | **Eiger** | **Wgn** | **Grnd** |
| *Actinia tenebrosa* | 7 | 20 | 0 |
| *Hydra vulgaris* | 5 | 1 | 0 |
| *Nematostella vectensis* | 4 | 11 | 0 |
| *Pocillopora damicornis* | 9 | 21 | 2 |
| *Stylophora pistillata* | 7 | 37 | 0 |
| **Total** | **32** | **90** | **2** |

| **PHYLUM PLATYHELMINTHES** | | | |
| --- | --- | --- | --- |
| **Species** | **Eiger** | **Wgn** | **Grnd** |
| *Macrostomum lignano* | 6 | 16 | 1 |
| *Schistosoma mansoni* | 0 | 3 | 0 |
| **Total** | **6** | **19** | **1** |

| **PHYLUM BRACHIOPODA** | | | |
| --- | --- | --- | --- |
| **Species** | **Eiger** | **Wgn** | **Grnd** |
| *Lingula unguis* | 17 | 11 | 6 |
| **Total** | **17** | **11** | **6** |

| **PHYLUM ANNELIDA** | | | |
| --- | --- | --- | --- |
| **Species** | **Eiger** | **Wgn** | **Grnd** |
| *Capitella teleta* | 9 | 2 | 12 |
| *Helobdella robusta* | 4 | 1 | 0 |
| *Dimorphilus gyrociliatus* | 9 | 2 | 2 |
| **Total** | **22** | **5** | **14** |

| **PHYLUM MOLLUSCA** | | | |
| --- | --- | --- | --- |
| **Species** | **Eiger** | **Wgn** | **Grnd** |
| *Crassostrea virginica* | 23 | 14 | 1 |
| *Octopus vulgaris* | 8 | 4 | 0 |
| *Mizuhopecten yessoensis* | 27 | 16 | 3 |
| *Lottia gigantea* | 5 | 10 | 0 |
| **Total** | **63** | **44** | **4** |

| **PHYLUM NEMATODA** | | | |
| --- | --- | --- | --- |
| **Species** | **Eiger** | **Wgn** | **Grnd** |
| *Caenorhabditis elegans* | 0 | 2 | 0 |
| *Onchocerca volvulus* | 0 | 0 | 0 |
| *Cercopithifilaria johnstoni* | 0 | 0 | 0 |
| *Loa loa* | 0 | 0 | 0 |
| *Brugia pahangi* | 0 | 0 | 0 |
| *Thelazia callipaeda* | 0 | 0 | 0 |
| *Litomosoides sigmodontis* | 0 | 0 | 0 |
| *Ancylostoma ceylanicum* | 0 | 2 | 0 |
| *Toxocara canis* | 0 | 1 | 0 |
| *Diploscapter pachys* | 0 | 0 | 0 |
| *Acanthocheilonema viteae* | 0 | 0 | 0 |
| **Total** | **0** | **5** | **0** |

| **PHYLUM TARDIGRADA** | | | |
| --- | --- | --- | --- |
| **Species** | **Eiger** | **Wgn** | **Grnd** |
| *Hypsibius exemplaris* | 0 | 3 | 0 |
| *Ramazzottius varieornatus* | 0 | 1 | 0 |
| **Total** | **0** | **4** | **0** |

| **PHYLUM ARTHROPODA** | | | |
| --- | --- | --- | --- |
| **SUBPHYLUM INSECTA** | | | |
| **Species** | **Eiger** | **Wgn** | **Grnd** |
| *Zootermopsis nevadensis* | 2 | 2 | 1 |
| *Coptotermes formosanus* | 0 | 2 | 1 |
| *Cryptotermes secundus* | 3 | 2 | 1 |
| *Ignelater luminosus* | 1 | 1 | 1 |
| *Photinus pyralis* | 1 | 0 | 1 |
| *Sitophilus oryzae* | 1 | 1 | 1 |
| *Tribolium castaneum* | 1 | 1 | 1 |
| *Rhynchophorus ferrugineus* | 1 | 1 | 1 |
| *Agrilus planipennis* | 1 | 1 | 1 |
| *Dendroctonus ponderosae* | 0 | 1 | 0 |
| *Drosophila melanogaster* | 1 | 1 | 1 |
| *Drosophila albomicans* | 1 | 1 | 1 |
| *Drosophila pseudoobscura* | 1 | 1 | 1 |
| *Drosophila lebanonensis* | 1 | 1 | 1 |
| *Drosophila ananassae* | 1 | 1 | 1 |
| *Aedes aegypti* | 1 | 1 | 1 |
| *Drosophila mojavensis* | 1 | 1 | 1 |
| *Drosophila erecta* | 1 | 1 | 1 |
| *Drosophila virilis* | 1 | 1 | 1 |
| *Anopheles gambiae* | 1 | 1 | 1 |
| *Musca domestica* | 1 | 1 | 1 |
| *Drosophila willistoni* | 1 | 1 | 1 |
| *Drosophila grimshawi* | 1 | 1 | 1 |
| *Drosophila kikkawai* | 1 | 1 | 1 |
| *Anopheles stephensi* | 1 | 1 | 1 |
| *Hermetia illucens* | 1 | 1 | 1 |
| *Polypedilum vanderplanki* | 1 | 2 | 1 |
| *Drosophila navojoa* | 1 | 0 | 1 |
| *Drosophila busckii* | 1 | 1 | 0 |
| *Drosophila sechellia* | 1 | 1 | 1 |
| *Anopheles darlingi* | 1 | 0 | 1 |
| *Clunio marinus* | 1 | 1 | 1 |
| *Culex quinquefasciatus* | 1 | 1 | 1 |
| *Drosophila rhopaloa* | 1 | 1 | 1 |
| *Lucilia cuprina* | 1 | 1 | 1 |
| *Drosophila persimilis* | 1 | 1 | 1 |
| *Anopheles sinensis* | 1 | 0 | 0 |
| *Cloeon dipterum* | 1 | 3 | 1 |
| *Acyrthosiphon pisum* | 4 | 1 | 1 |
| *Laodelphax striatellus* | 2 | 4 | 1 |
| *Sipha flava* | 2 | 1 | 1 |
| *Cimex lectularius* | 2 | 2 | 1 |
| *Apolygus lucorum* | 2 | 0 | 1 |
| *Cinara cedri* | 2 | 1 | 3 |
| *Rhodnius prolixus* | 1 | 1 | 0 |
| *Aphis craccivora* | 0 | 1 | 1 |
| *Diaphorina citri* | 2 | 1 | 1 |
| *Pogonomyrmex barbatus* | 1 | 3 | 1 |
| *Cyphomyrmex costatus* | 1 | 1 | 1 |
| *Temnothorax curvispinosus* | 1 | 2 | 1 |
| *Trachymyrmex septentrionalis* | 1 | 1 | 1 |
| *Atta cephalotes* | 1 | 2 | 1 |
| *Trachymyrmex cornetzi* | 1 | 1 | 0 |
| *Ooceraea biroi* | 2 | 2 | 1 |
| *Apis mellifera* | 2 | 2 | 1 |
| *Apis cerana* | 2 | 2 | 0 |
| *Bombus vosnesenskii* | 2 | 2 | 1 |
| *Dinoponera quadriceps* | 1 | 2 | 1 |
| *Vespula pensylvanica* | 1 | 2 | 1 |
| *Fopius arisanus* | 1 | 2 | 1 |
| *Nasonia vitripennis* | 0 | 1 | 0 |
| *Atta colombica* | 1 | 2 | 1 |
| *Habropoda laboriosa* | 2 | 2 | 1 |
| *Camponotus floridanus* | 2 | 2 | 1 |
| *Frieseomelitta varia* | 2 | 2 | 1 |
| *Vespula germanica* | 1 | 2 | 0 |
| *Trachymyrmex zeteki* | 1 | 2 | 0 |
| *Acromyrmex echinatior* | 1 | 2 | 1 |
| *Aphidius gifuensis* | 2 | 2 | 1 |
| *Acromyrmex charruanus* | 0 | 0 | 0 |
| *Harpegnathos saltator* | 1 | 1 | 1 |
| *Melipona quadrifasciata* | 2 | 1 | 0 |
| *Neodiprion lecontei* | 2 | 2 | 1 |
| *Dufourea novaeangliae* | 2 | 1 | 1 |
| *Spodoptera litura* | 2 | 2 | 1 |
| *Vanessa tameamea* | 2 | 1 | 1 |
| *Bombyx mori* | 3 | 3 | 1 |
| *Papilio machaon* | 1 | 2 | 1 |
| *Parnassius apollo* | 2 | 2 | 1 |
| *Trichoplusia ni* | 2 | 3 | 1 |
| *Arctia plantaginis* | 2 | 3 | 1 |
| *Danaus plexippus* | 2 | 1 | 1 |
| *Danaus chrysippus* | 2 | 1 | 0 |
| *Pieris macdunnoughi* | 1 | 1 | 1 |
| *Papilio xuthus* | 2 | 2 | 1 |
| *Brenthis ino* | 0 | 0 | 0 |
| *Manduca Sexta* | 2 | 4 | 1 |
| *Leptidea sinapis* | 2 | 1 | 1 |
| *Plutella xylostella* | 2 | 1 | 1 |
| *Heliothis virescens* | 2 | 2 | 0 |
| *Pararge aegeria aegeria* | 2 | 1 | 0 |
| *Operophtera brumata* | 2 | 0 | 0 |
| *Ladona fulva* | 1 | 2 | 0 |
| *Frankliniella occidentalis* | 3 | 0 | 0 |
| *Thrips palmi* | 2 | 2 | 1 |
| *Allacma fusca* | 2 | 2 | 1 |
| *Orchesella cincta* | 3 | 0 | 2 |
| **Total** | **136** | **134** | **85** |

| **SUBPHYLUM CRUSTACEA** | | | |
| --- | --- | --- | --- |
| **Species** | **Eiger** | **Wgn** | **Grnd** |
| *Darwinula stevensoni* | 2 | 1 | 1 |
| *Hyalella azteca* | 1 | 0 | 0 |
| *Daphnia pulex* | 1 | 2 | 0 |
| *Daphnia galeata* | 2 | 3 | 0 |
| *Daphnia magna* | 1 | 1 | 0 |
| *Tigriopus californicus* | 1 | 1 | 0 |
| *Homarus americanus* | 1 | 0 | 0 |
| *Amphibalanus amphitrite* | 1 | 1 | 2 |
| **Total** | **10** | **9** | **3** |

| **SUBPHYLUM MYRIAPODA** | | | |
| --- | --- | --- | --- |
| **Species** | **Eiger** | **Wgn** | **Grnd** |
| *Strigamia maritima* | 2 | 1 | 0 |
| **Total** | **2** | **1** | **0** |

| **SUBPHYLUM CHELICERATA** | | | |
| --- | --- | --- | --- |
| **Species** | **Eiger** | **Wgn** | **Grnd** |
| *Varroa destructor* | 0 | 0 | 0 |
| *Trichonephila clavata* | 0 | 2 | 0 |
| *Dermatophagoides pteronyssinus* | 0 | 1 | 0 |
| *Dermatophagoides farinae* | 0 | 1 | 0 |
| *Sarcoptes scabiei* | 0 | 1 | 0 |
| *Araneus ventricosus* | 0 | 2 | 0 |
| *Tetranychus urticae* | 0 | 0 | 0 |
| *Argiope bruennichi* | 0 | 2 | 0 |
| *Dinothrombium tinctorium* | 0 | 2 | 0 |
| *Stegodyphus mimosarum* | 0 | 0 | 0 |
| *Ixodes scapularis* | 0 | 0 | 0 |
| *Leptotrombidium deliense* | 0 | 0 | 0 |
| **Total** | **0** | **11** | **0** |

**Supplementary Table S4: Number of Eiger, wengen and grnd homologs per species after searching UniPtot database using in-house HMM model.**

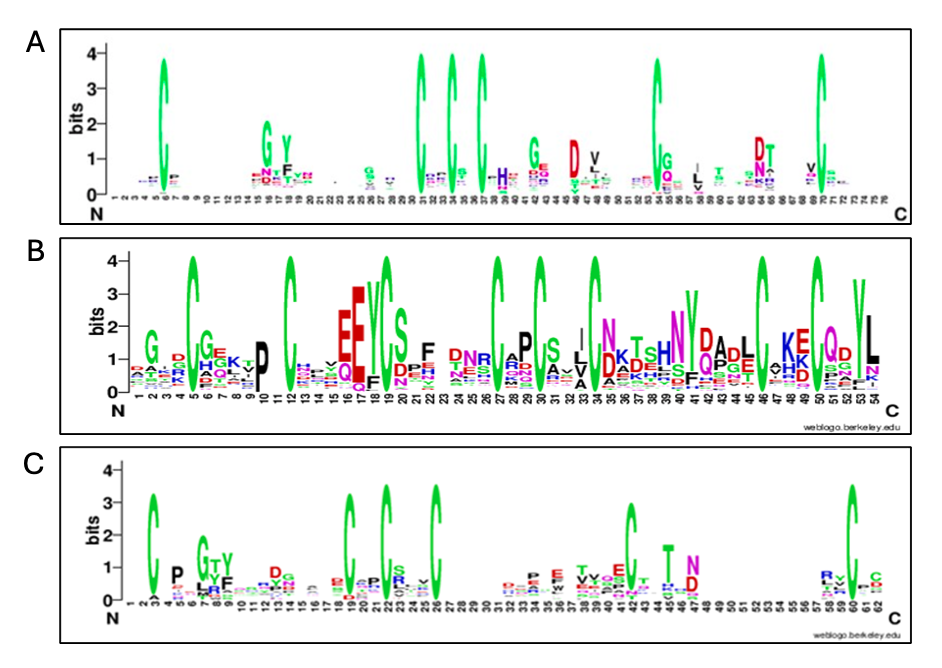

**Supplementary Figure S1(a):** The seed alignment of the Pfam TNFR model (PF00020.21) is presented as a sequence logo with 489 TNFR protein sequences containing CXXCXXC THDs which is 71 amino acids long. **(b)** The arthropod grindelwald seed alignment collected in-house is presented as a sequence logo 79 Grnd protein sequences containing CXXCXXXC domains which are ~54 amino acids in length. **(c)** The seed alignment of the invertebrate Grnd model is represented as a sequence logo with 21 Grnd homologs containing THDs which is 60 amino acids long.

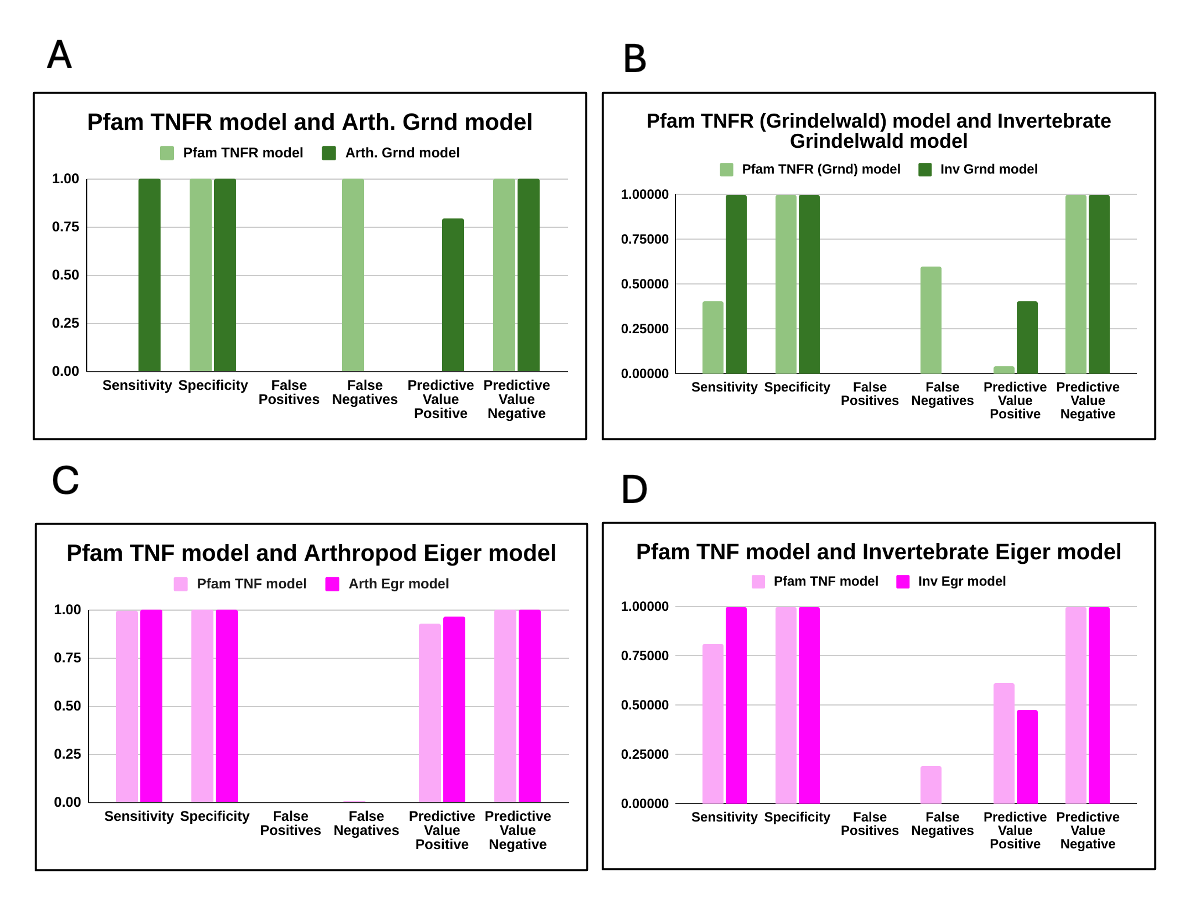

**Supplementary Figure S2:** Bar plots comparing HMM models for TNF and TNFR domains in Pfam with those developed in-house. a) TNFR (PF0020.21) and the Arthropod Grnd model built in-house. **(b)** TNFR model (PF00020.21) and the Invertebrate Grnd model built in-house.

- Sensitivity = #TNF/TNFR Positive Hits / #Total TNF-TNFR Hits
- Specificity = #Non-TNF/TNFR Negative Hits / #Total Non-TNF-TNFR Hits
- False Positives = #Non-TNF/TNFR Positive Hits / #Total Non-TNF /TNFR Hits
- False Negatives = # TNF/TNFR Positive Hits/ # Total TNF/TNFR Hits
- Predictive Value Positive = #Total Positive TNF/TNFR Hits/ #Total Positive TNF/TNFR + Non TNF/TNFR Hits
- Predictive Value Negative = #Non-TNF/TNFR Negative Hits/ #Total Negative TNF/TNFR + Non TNF/TNFR Hits

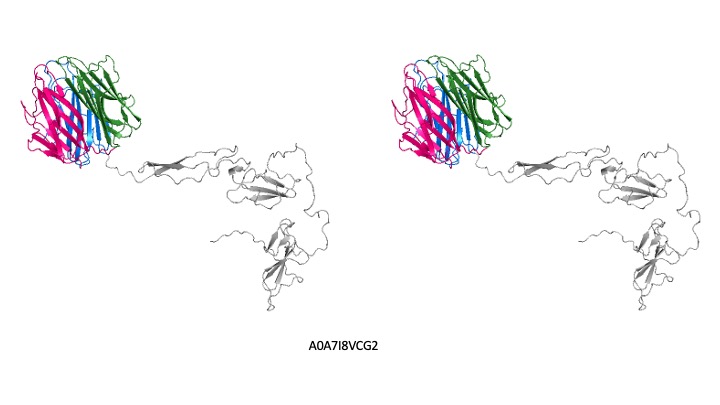

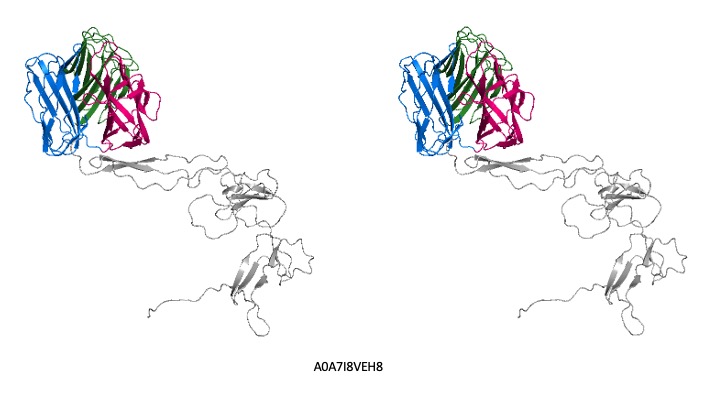

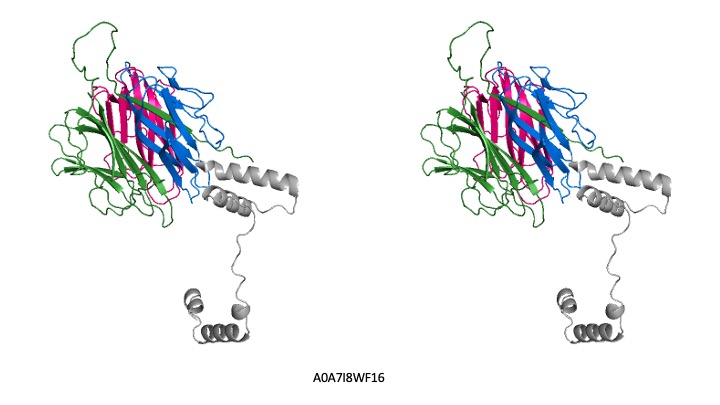

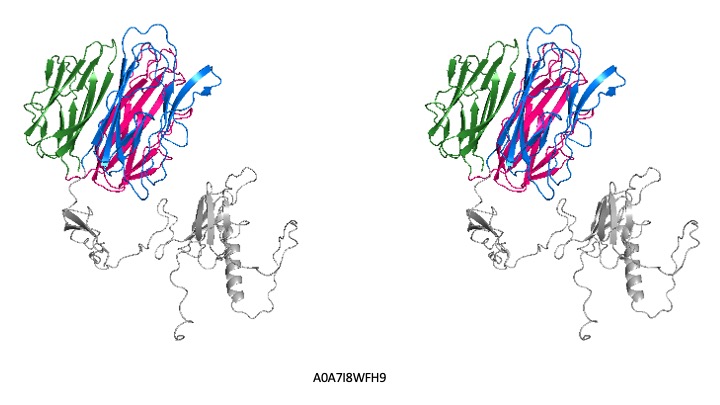

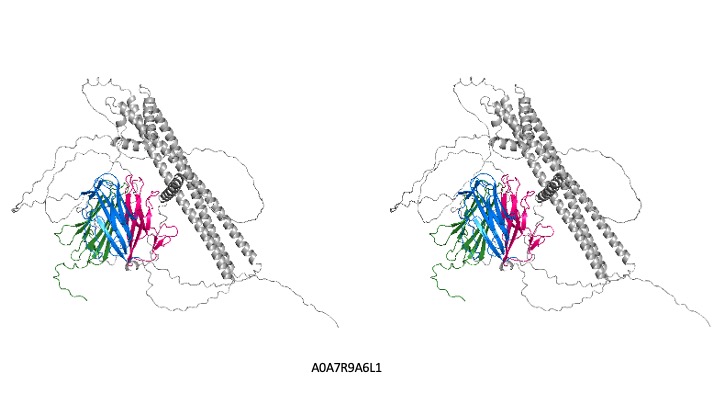

Supplementary Figure S3: Stereo of all the molecules shown in Figure 3.

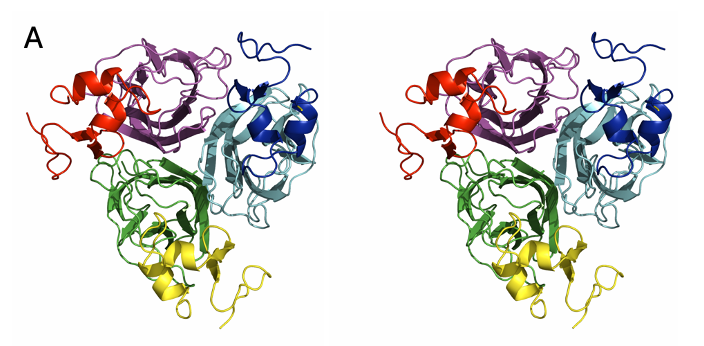

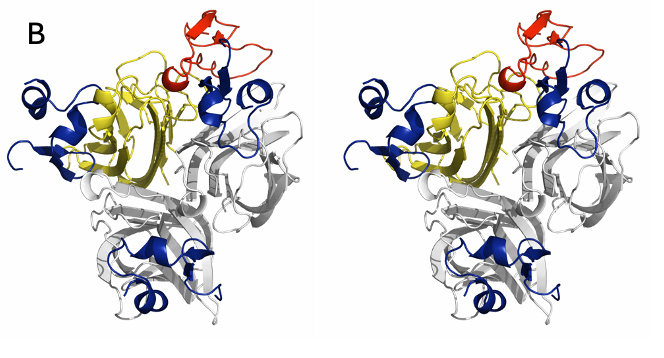

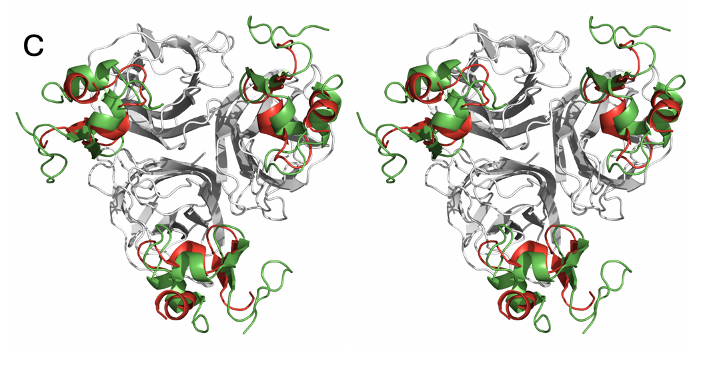

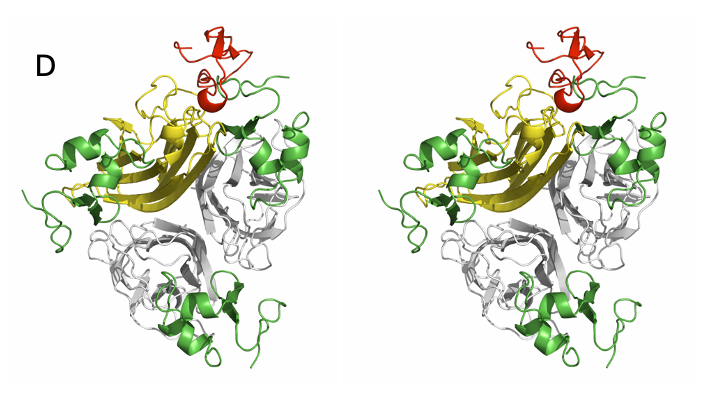

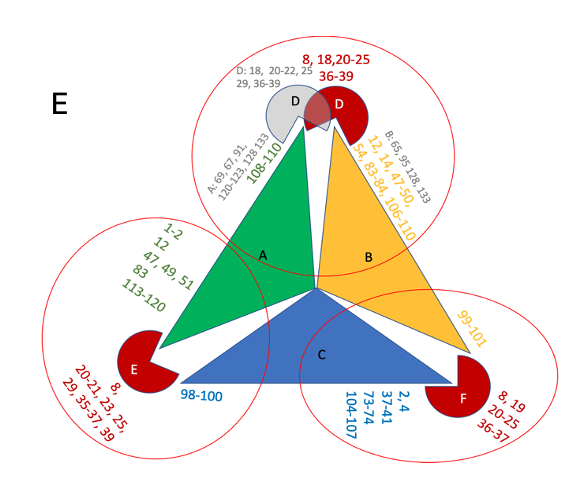

**Supplementary Figure S4(a):** Stereo view of the structure of the complex of homotrimer of DmEiger from 6zt0 (green, cyan and violet) along with three copies of its receptor binding domain (red, yellow and blue). **(b)** 1TNR crystal structure (Yellow - ligand and Red - -receptor) superimposed on APRIL-BCMA complex (1XU2). **(c)** APRIL- BCMA (Red) superimposed on 6ZT0 (green - receptor, grey - ligand). **(d)**  1TNR (Yellow - ligand, Red - receptor) superimposed on 6ZT0 (Green - receptor, Grey - ligand). **(e)** residue-residue contact of ligand-receptor chains from 6ZT0 and complex of dsEiger-DsGrnd shown in Figure 5a.

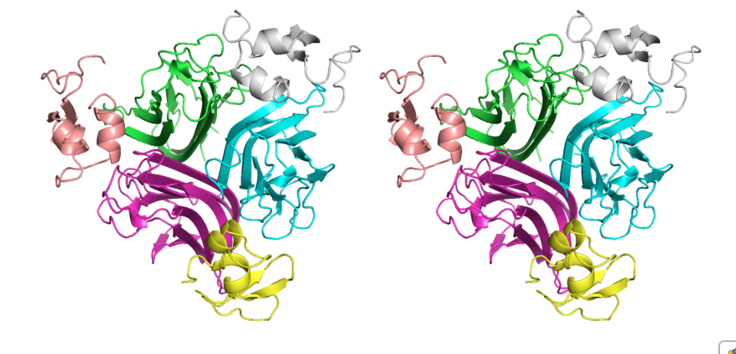

**Supplementary Figure S5:** Stereo view of AlphaFold model of HuTNF-alpha-DmGrnd complex -built as a negative control.

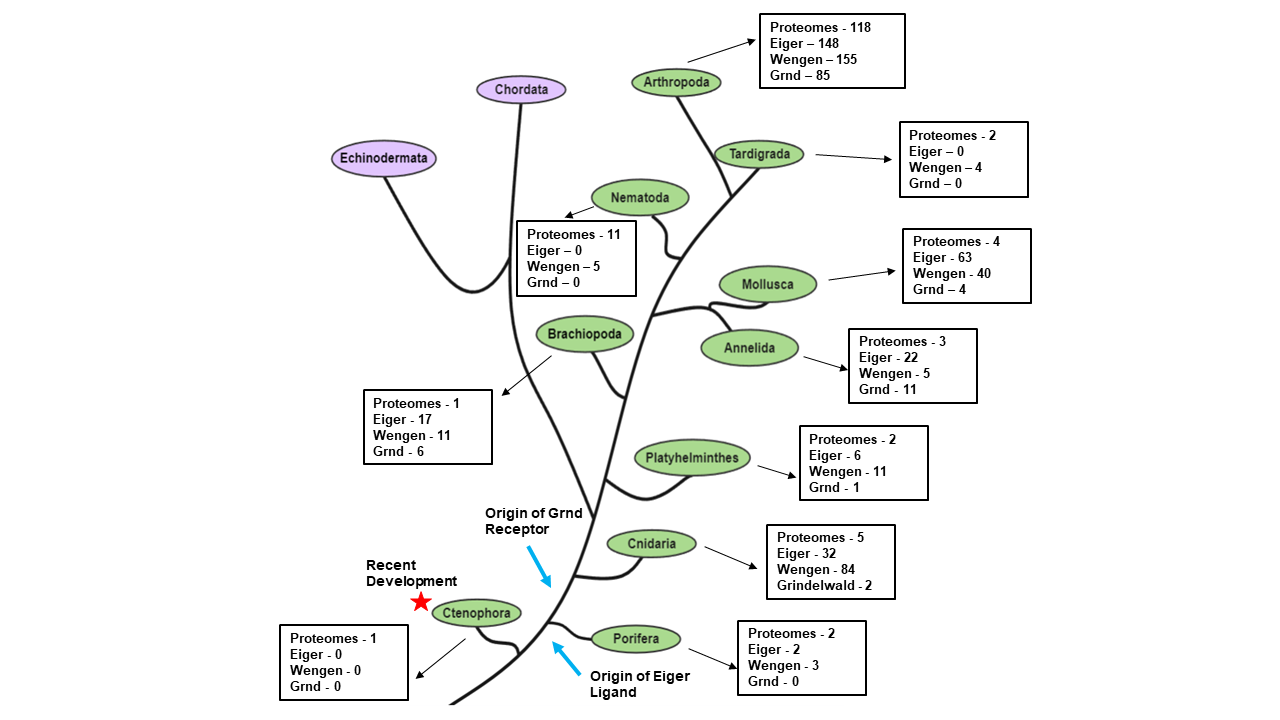

Supplementary Figure S6: Tree of metazoa showing the number of proteomes mined, and number of Eiger, wgn and grnd homologs identified under each phyla.

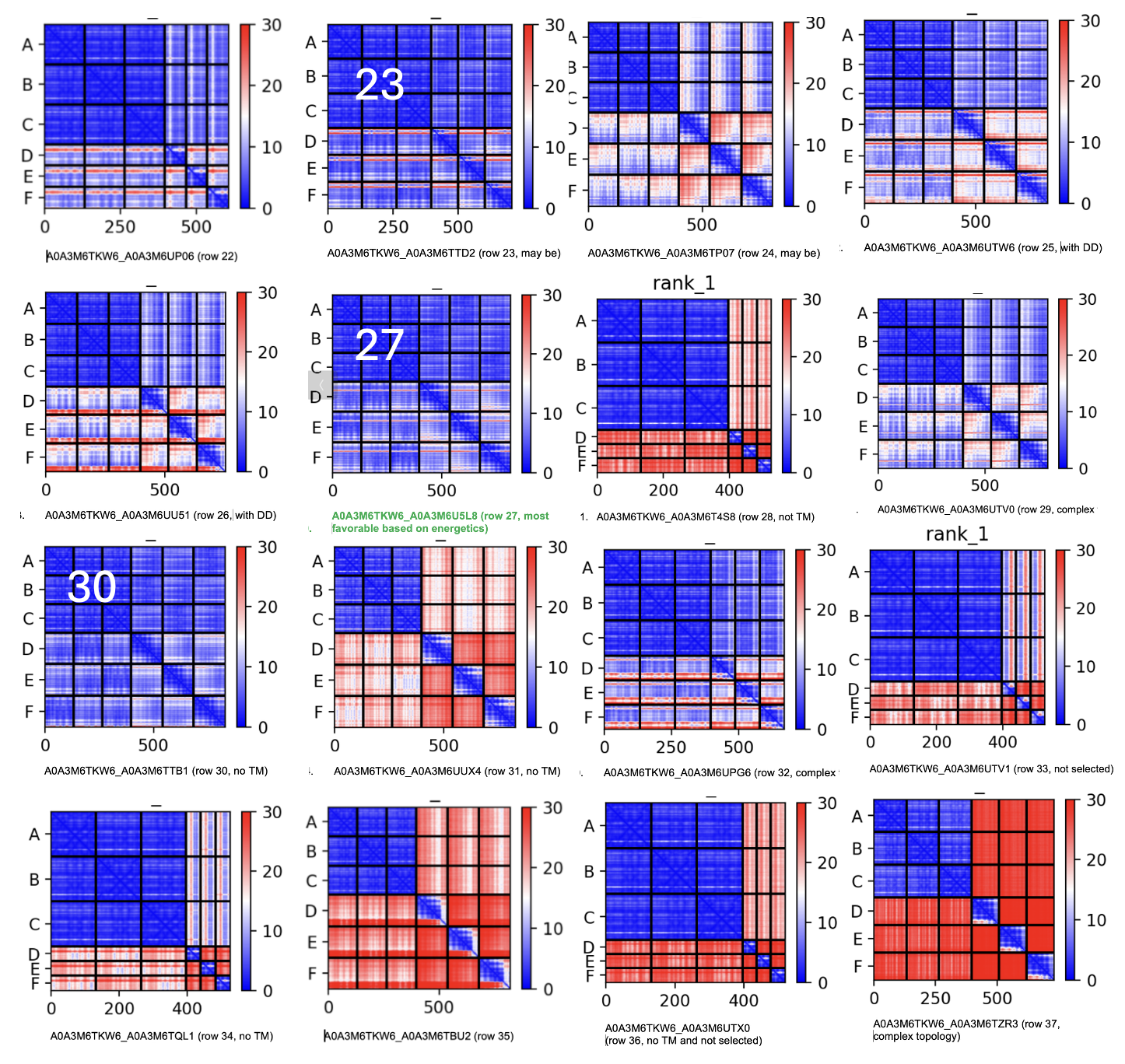

Supplementary Figure S7: Amino acid-Amino acid predicted align error (PAE) for experiment spanning rows 22-37 in Table 1. The most confident models based with low PAE align are rows 23, 27 and 30 of Table 1, with row 27 having the lowest delta-G and satisfying other functional requirement such as presence of TM etc as shown in Table 1.

| Sr | ID | Mean pae | deltaG | Functional/strctural complexity |
| --- | --- | --- | --- | --- |
| 22 | A0A3M6TKW6_A0A3M6UP06 | 7.425 | -21.2 | fail interface |
| 23 | A0A3M6TKW6_A0A3M6TTD2 | 6.770 | -27 | OK |
| 24 | A0A3M6TKW6_A0A3M6TP07 | 10.590 | -27 | High pae |
| 25 | A0A3M6TKW6_A0A3M6UTW6 | 9.871 | -18.9 | OK |
| 26 | A0A3M6TKW6_A0A3M6UU51 | 10.426 | -17.9 | Low deltaG high pae |
| **27** | **A0A3M6TKW6_A0A3M6U5L8** | **6.753** | **-35** | **BEST** |
| 28 | A0A3M6TKW6_A0A3M6T4S8 | 14.089 | -19.2 | High pae |
| 29 | A0A3M6TKW6_A0A3M6UTV0 | 8.810 | -20 | High pae |
| 30 | A0A3M6TKW6_A0A3M6TTB1 | 6.786 | .30.5 | OK |
| 31 | A0A3M6TKW6_A0A3M6UUX4 | 15.173 | -20.6 | High Pae |
| 32 | A0A3M6TKW6_A0A3M6UPG6 | 8.742 | -24.9 | Complex topology |
| 33 | A0A3M6TKW6_A0A3M6UTV1 | 9.945 | -16 | Low deltaG |
| 34 | A0A3M6TKW6_A0A3M6TQL1 | 17.325 | -23.3 | High pae |
| 35 | A0A3M6TKW6_A0A3M6TBU2 | 17.560 | -10.6 | High pae |
| 36 | A0A3M6TKW6_A0A3M6UTX0 | 11.722 | -26.7 | High pae |
| 37 | A0A3M6TKW6_A0A3M6TZR3 | 19.152 | -11.6 | High pae |

Supplementary Table S5: The table provides predicted aligned error (column 3) and respective deltaG from Table 1 for experiments listed in Table 1 from rows 22-37. Receptor in rows 23, 27 and 30 are potential receptors for A0A3M6TKW6 with A0A3M6U5L8 with lowest PAE and delatG
